# Isolating salient variations of interest in single-cell data with contrastiveVI

**DOI:** 10.1101/2021.12.21.473757

**Authors:** Ethan Weinberger, Chris Lin, Su-In Lee

## Abstract

Single-cell datasets are routinely collected to investigate changes in cellular state between control cells and corresponding cells in a treatment condition, such as exposure to a drug or infection by a pathogen. To better understand heterogeneity in treatment response, it is desirable to disentangle latent structures and variations uniquely enriched in treated cells from those shared with controls. However, standard computational models of single-cell data are not designed to explicitly separate these variations. Here, we introduce Contrastive Variational Inference (contrastiveVI; https://github.com/suinleelab/contrastiveVI), a framework for analyzing treatment-control scRNA-seq datasets that explicitly disentangles the data into shared and treatment-specific latent variables. Using four treatment-control scRNA-seq dataset pairs, we apply contrastiveVI to perform a broad set of standard analysis tasks, including visualization, clustering, and differential expression testing. In each case, we find that our method consistently achieves results that agree with known biological ground truths, while previously proposed methods often fail to do so. We conclude by generalizing our framework to multimodal measurements and applying it to analyze a single-cell dataset with joint transcriptome and surface protein measurements.

## Main

Single-cell technologies have emerged as powerful tools for understanding previously unexplored biological diversity. To facilitate the investigation of various biological hypotheses, single-cell data are often collected simultaneously from cells in a treatment condition and from control cells. For example, recent studies have profiled cells from tumors versus healthy tissue [1], cells exposed to drug compounds versus placebos [2], and cells with CRISPR-induced genomic perturbations versus cells with unaltered genomes [3, 4]. To better understand a given phenomenon under investigation, it is desirable to isolate the low-dimensional structures and variations enriched in data from the *target* cells (i.e., cells in the treatment condition) and compare them to a corresponding set of *background* cells (i.e., cells in the control condition). With the development of new technologies for measuring cellular responses to large numbers of perturbations in parallel, such as Perturb-Seq [3], MIX-Seq [2], and MULTI-Seq [1] among others, tools for a refined understanding of variations unique to target vs. background cells will be increasingly critical.

Isolating the variations enriched in a target dataset is the subject of *contrastive analysis* (CA) [5, 6, 7, 8, 9, 10]. Many recent studies proposed probabilistic latent variable models for analyzing single-cell data [11, 12, 13]. However, these methods are not suitable for CA since they use a *single* set of latent variables to model *all* of the variations in the data. Because the variations specifically enriched in a target dataset are often subtle compared to the overall variations in the data [6], such models will likely strongly entangle the enriched variations of interest with irrelevant latent factors or fail to capture the enriched variations entirely.

We note the fundamental difference in the goals between CA and batch effect correction, which has been extensively studied in previous work [14, 15, 16]. When correcting for batch effects between datasets, it is assumed that systematic variations between datasets are due to irrelevant technical factors and should be discarded. On the other hand, in the CA setting we assume that variations enriched in the target dataset are the result of meaningful biological phenomena and should be isolated for further study. This assumption is reasonable under many common experimental designs (**Supplementary Note 1**) and is borne out in our empirical results. We also emphasize that the goal of CA is distinct from the related problem of differential abundance testing [17, 18, 19]. Methods for differential abundance testing aim to identify subpopulations of cells whose distributions differ across a target and background state. However, these methods do not provide any mechanisms for understanding how identified populations of cells relate to each other. In contrast, by isolating the variations enriched in target cells from those shared with background cells, CA methods may facilitate a more comprehensive understanding of the relationships between populations of target cells, such as activation of a similar biological mechanism. We provide a more detailed overview of the differences between CA and differential abundance testing in **Supplementary Note 2**.

Despite the many potential use cases for CA methods with single-cell data, little work has been done on the subject. One recent study [20] proposed a nearest-neighbors-based algorithm, called mixscape, for removing confounding variations shared with control cells from CRISPR-perturbed single-cell transcriptome measurements. However, this algorithm operates on normalized count data rather than raw counts. This property is not ideal, since many normalization approaches have been proposed with no single method being appropriate for all types of scRNA-seq data [21], and multiple studies [12, 22] have found that choice of normalization procedure substantially impacts downstream results. Moreover, once mixscape is applied, other tools are needed to perform further analysis tasks. Previous work [11, 23] has advocated against such workflows, arguing instead that using a single distributional model for a wider range of analysis tasks yields more consistent and interpretable results. Another recent work [7] designed a probabilistic model for analyzing raw scRNA-seq count data in the CA setting. Though this method has provided new insights into variations enriched in target scRNA-seq datasets, it assumes that a linear model is sufficiently expressive to model scRNA-seq data, despite previous work that demonstrated substantial improvements by using more expressive nonlinear methods [11]. Furthermore, none of these previously proposed methods can incorporate information from other modalities beyond scRNA-seq.

To address these limitations, we developed Contrastive Variational Inference (contrastiveVI), a deep generative model for analyzing scRNA-seq data in the CA setting (**Fig. 1)**. contrastiveVI models the variations underlying scRNA-seq data using two sets of latent variables: the first, called *shared variables*, are common to background and target cells, while the second, called *salient variables*, are used to model variations specific to target data. Similar to previous work [11], our probabilistic model accounts for the specific technical biases and noise characteristics of scRNA-seq data. To our knowledge, contrastiveVI is the first probabilistic model designed for analyzing single-cell data in the CA setting that can capture the complex nonlinear variations present in scRNA-seq data. contrastiveVI can be used for many analysis tasks, including dimensionality reduction, clustering, and differential gene expression testing. To highlight this functionality, we applied contrastiveVI to four publicly available background and target scRNA-seq dataset pairs. We found that contrastiveVI consistently achieved results that agree with known biological ground truths. We also generalized our framework to multimodal datasets by developing totalContrastiveVI, a model for analyzing joint RNA and surface protein measurements in the CA setting, and applied it to analyze CRISPR-induced variations in an ECCITE-Seq [24] dataset.

**Figure 1:**
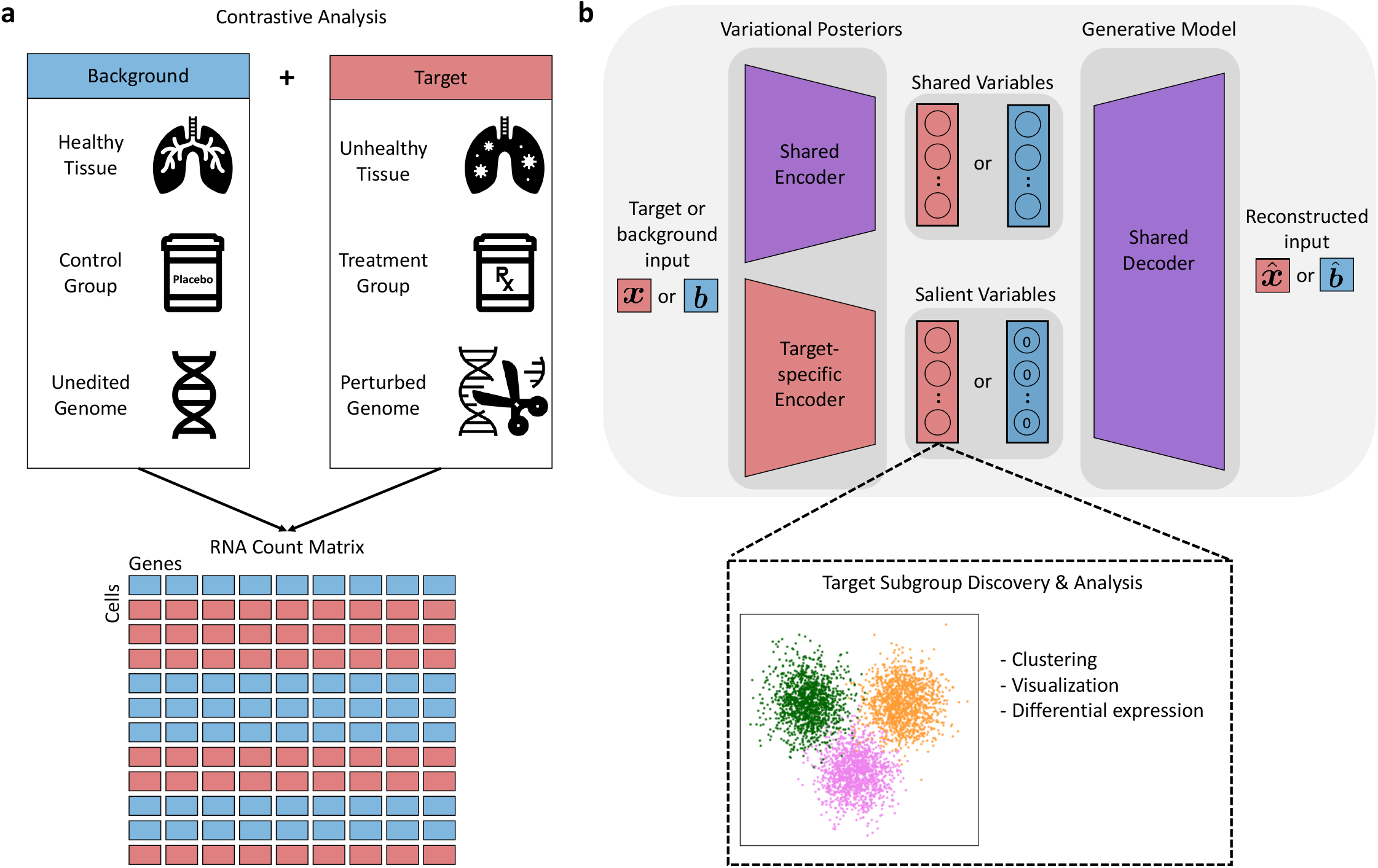
Overview of contrastiveVI. Given a reference background dataset and a target dataset of interest, contrastiveVI separates the variations shared between the two datasets and the variations enriched in the target dataset. **a**, Example background and target data pairs. Samples from both conditions produce an RNA count matrix with each cell labeled as background or target. **b**, Schematic of the contrastiveVI model. A shared encoder network embeds a cell, whether target (red) or background (blue), into the model’s shared latent space, which captures variations common to target and background cells. A second target-cell-specific encoder embeds target cells into the model’s salient latent space, which captures variations enriched in the target data and not present in the background. For background cells the values of the salient latent factors are fixed to be a zero vector. Both target and background cells’ latent representations are transformed back to the original gene expression space using a single shared decoder network.

## Results

### The contrastiveVI model

contrastiveVI uses a probabilistic latent variable model to represent the uncertainty in observed RNA counts as a combination of biological and technical factors. Model input consists of an RNA count matrix and labels denoting whether each cell is a background or target cell (**Fig. 1a**). Additional categorical covariates, such as anonymized donor ID or experimental batch, are optional inputs to the model that can be used to integrate datasets.

contrastiveVI encodes each cell as the parameters of a distribution in a low-dimensional latent space. This latent space is divided into two parts, each with its own encoding function. The first set of latent variables, called *shared variables*, captures factors of variation common to both background and target data. The second set, denoted as *salient variables*, captures variations unique to the target dataset. Only target data points are assigned salient latent variable values; background data points are instead assigned a zero vector for the salient variables to represent their absence. contrastiveVI also provides a way to estimate the parameters of the distributions underlying the observed RNA measurements given a cell’s latent representation. Such distributions explicitly account for technical factors in the observed data, such as sequencing depth, batch effects, and dropouts. All distributions are parameterized by neural networks.

The contrastiveVI model is based on the variational autoencoder (VAE) framework [25]. As such, its parameters can be learned using efficient stochastic optimization techniques, easily scaling to large scRNA-seq datasets consisting of tens or hundreds of thousands of cells. Following optimization, we can make use of the various contrastiveVI model components for downstream analyses. For example, the salient latent representations of target samples can be used as inputs to clustering or visualization algorithms to discover subgroups of target points. Moreover, the distributional parameters can be used for additional tasks such as imputation or differential gene expression analysis. A more detailed description of the contrastiveVI model can be found in **Methods**.

### contrastiveVI isolates subtle variations in target cells

To evaluate the performance of contrastiveVI and other methods, we rely on datasets with known biological variations in the target condition that are not present in the background condition. One such dataset consists of expression data from bone marrow mononuclear cells (BMMCs) from two patients with acute myeloid leukemia (AML) and two healthy controls [26]. The two patients underwent allogenic stem-cell transplants, and BMMC samples were collected pre- and post-transplant. It is known that gene expression profiles of BMMCs differ pre- and post-transplant [26]. Therefore, the known biological variations in this target dataset (AML patient BMMCs) correspond to pre- vs. post-transplant cellular states, and a performant model should learn a salient latent space that differentiates pre- vs. post-transplant cells. We find qualitatively that pre- and post-transplant cells are indeed well separated in the salient latent space learned by contrastiveVI when trained using data from the healthy patients as the background dataset (**Fig. 2a**).

**Figure 2:**
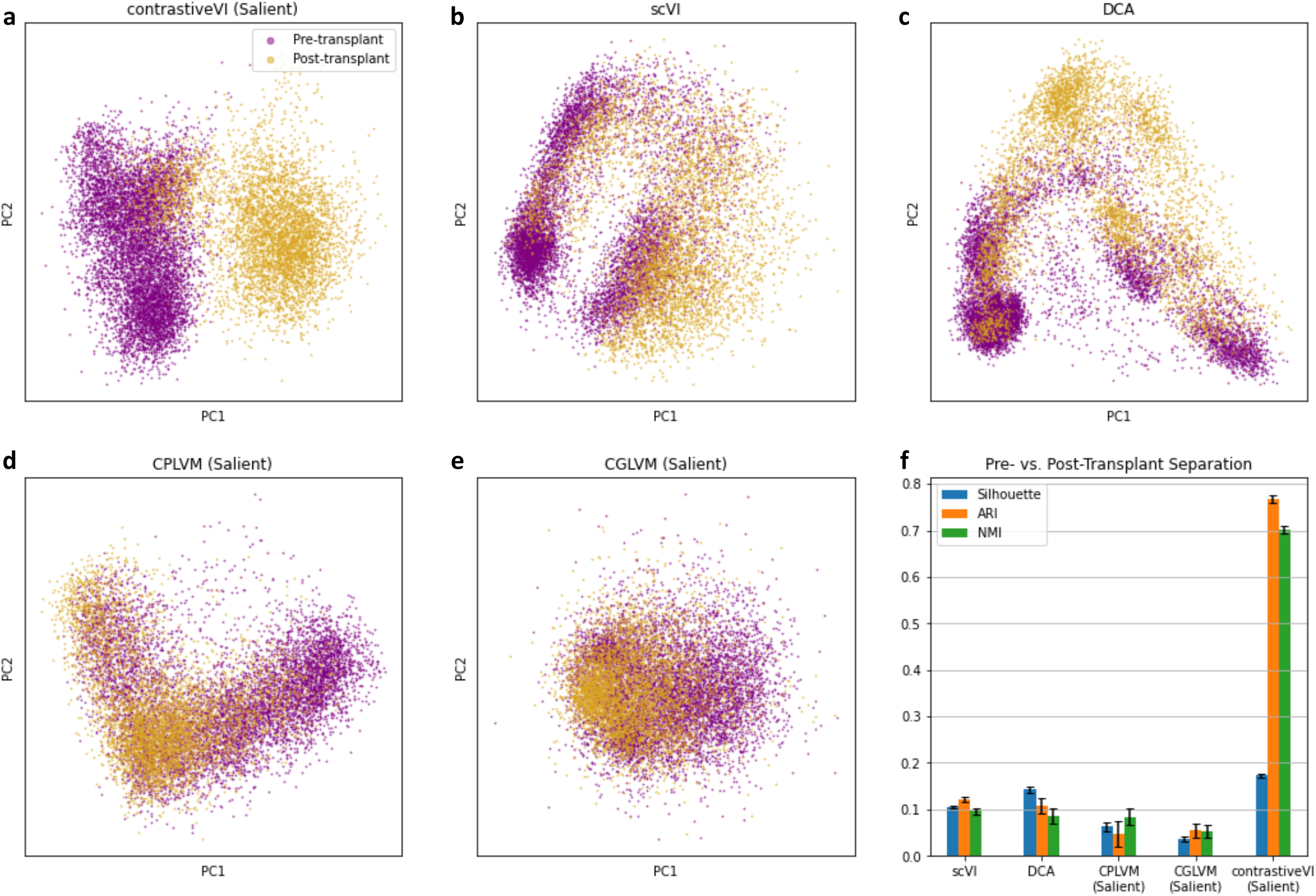
contrastiveVI successfully captures enriched variations in scRNA-seq data. **a-e**, Principal component (PC) plots of contrastiveVI and baseline models’ latent representations. For scVI and DCA, the first two PCs of the models’ single latent representations are plotted, while for contrastive methods the PCs from their salient latent representations are plotted. **f**, Quantitative measures of separation between pre- and post-transplant cells. Silhouette is the average silhouette width of pre- vs. post-transplant cells, ARI is the adjusted Rand index, and NMI is the normalized mutual information. Higher values indicate better performance for all metrics. For each method, the mean and standard error across five random trials are plotted.

We compared contrastiveVI’s performance against that of four previously proposed methods for analyzing scRNA-seq data. Because the choice of library size normalization method has been shown to drastically impact dimension reduction and subsequent clustering results of methods not designed to explicitly model library sizes [12], we restricted our attention to methods that operate directly on unnormalized scRNA-seq count data. First, to demonstrate that our contrastive approach is necessary for isolating enriched variations in target datasets, we compared against single-cell variational inference (scVI) [11] and the deep count autoencoder (DCA) [27]. scVI and DCA have been successfully applied to perform a variety of scRNA-seq analysis tasks; however, they were not specifically designed for the CA setting and thus may struggle to capture salient variations in target samples. We also compared against two variants of the linear contrastive method designed for analyzing scRNA-seq count data proposed by Jones et al. [7]: contrastive Poisson latent variable model (CPLVM) and contrastive generalized latent variable model (CGLVM). Qualitatively (**Fig. 2b-e**), we find that none of these baseline models separate pre- and post-transplant cells as clearly as contrastiveVI.

We also quantified how well each method separates the two groups of target cells using three metrics— the average silhouette width, adjusted Rand Index (ARI), and normalized mutual information (NMI; **Methods**). Across all of our metrics, we find that contrastiveVI significantly outperforms baseline models (**Fig. 2f**), with especially large gains in the ARI and NMI. We provide the exact numerical values for these metrics in **Supplementary Table 1**. These results further indicate that contrastiveVI recovered the variations enriched in the AML patient data far better than baseline models.

Furthermore, we experimented with a workflow for using contrastiveVI for end-to-end biological discovery. After embedding the AML patient samples into the contrastiveVI salient latent space, we used k-means clustering to divide the target samples into two groups. Highly differentially expressed genes across the two clusters of target cells were then obtained by Monte Carlo sampling of denoised, library-size-normalized expressions from the contrastiveVI decoder to estimate a distribution over the log fold change of expression between the two sets of cells (**Methods**). The results of this differential expression test were validated by comparing the Bayes factors obtained from contrastiveVI with those obtained from scVI, which has previously been evaluated [11], when given the same clustering labels; we found the results from both tests were strongly correlated (Spearman’s *ρ* = 0.998 *±* 8.79 · 10^−5^). Finally, pathway enrichment analysis (**Methods**) was performed with these differentially expressed genes using the Kyoto Encyclopedia of Genes and Genomes (KEGG) 2016 pathway database [28]. Based on our quantitative results, our two clusters exhibited strong agreement with the two ground-truth groups, i.e., pre- and post-transplant cells. Moreover, as we might expect, the pathways enriched by the differentially expressed genes between the two clusters are related to immune response and graft rejection (**Supplementary Fig. 1)**. We provide a full list of enriched pathways in **Supplementary Table 2**. These results illustrate how contrastiveVI can enrich our understanding of variations specific to target scRNA-seq datasets.

### Separating infected mouse intestinal epithelial cells by pathogen type

We next applied contrastiveVI to data collected in Haber et al. [29]. This dataset consists of gene expression measurements of intestinal epithelial cells from mice infected with either *Salmonella enterica* (*Salmonella*) or *Heligmosomoides polygyrus* (*H. poly*). For the background dataset, we used measurements collected from healthy cells released by the same authors. Here, our goal is to separate cells by infection type in the contrastiveVI salient latent space.

We find that contrastiveVI successfully separates cells by infection type in its salient latent space (**Fig. 3a**). Moreover, we find that cells mix across infection types in the contrastiveVI shared latent space, as expected (**Fig. 3b**). These results indicate that enriched variations due to infection response are correctly relegated to the salient latent space. On the other hand, qualitatively (**Supplementary Fig. 2)** scVI and DCA strongly entangle shared and target-specific variations in their single latent spaces, while CPLVM and CGLVM do not clearly stratify the two classes of target samples in their salient latent spaces. These results were also reflected in our quantitative metrics (**Fig. 3c**).

**Figure 3:**
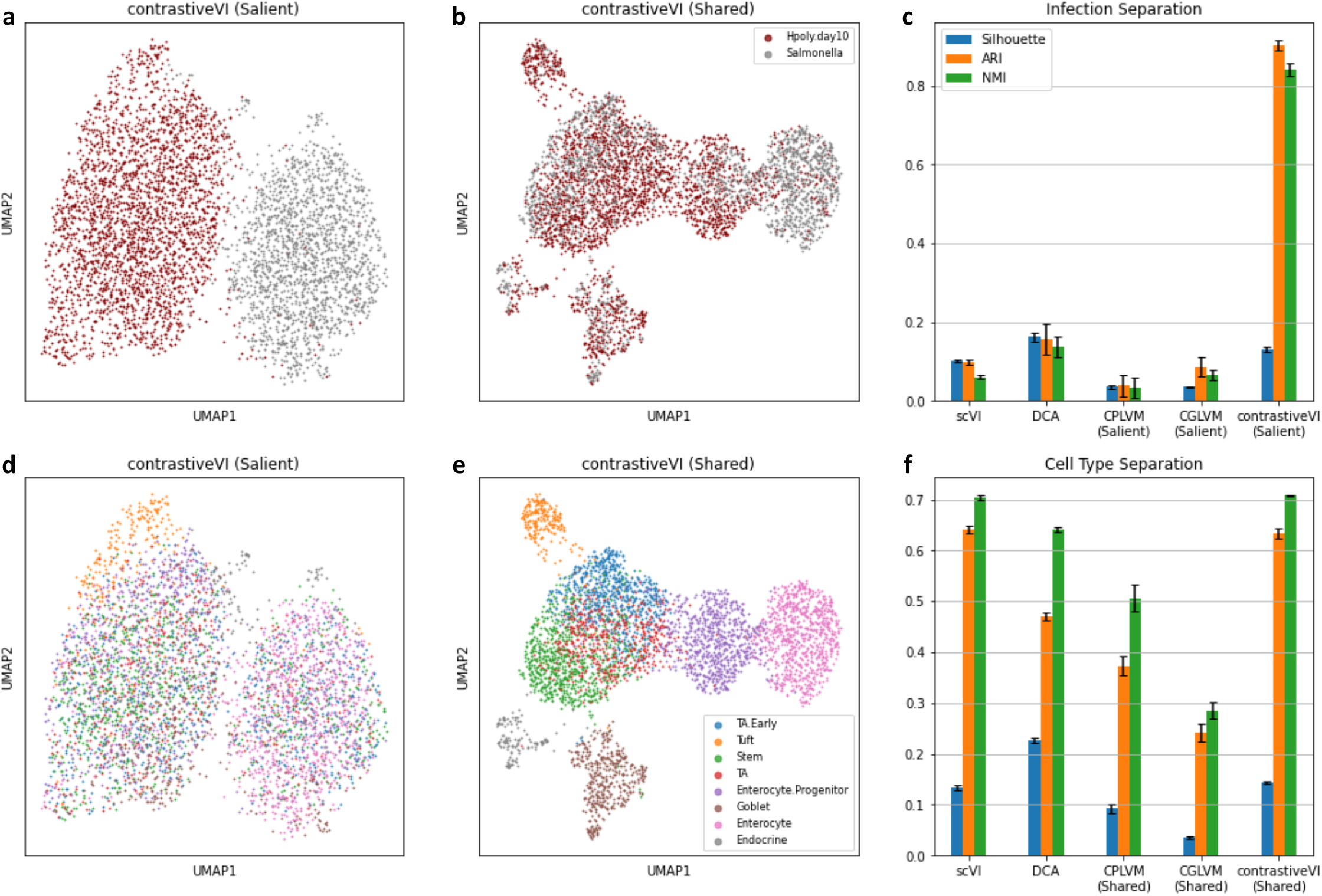
contrastiveVI isolates responses to different infections in mouse intestinal epithelial cells. **a**,**b**, UMAP plots of contrastiveVI’s salient and shared representations colored by infection type. Cells are correctly separated by infection type in the salient space, while they mix across infection types in the shared space. **c**, Clustering metrics quantify how well cells separate by infection type for scVI and DCA’s single latent spaces and contrastive models’ salient latent spaces, with means and standard errors across five random trials plotted. **d**,**e**, UMAP plots of contrastiveVI’s salient and shared representations colored by cell type. Cells separate well by cell type in the shared latent space, while they mix across cell types in the salient space. **f**, Quantifying how well cells separate by cell type in scVI and DCA’s single latent spaces and contrastive models’ shared latent spaces.

For this dataset we also validated contrastiveVI’s ability to separate salient and shared variations using ground truth cell type labels provided by Haber et al. [29]. We found strong mixing across cell types in contrastiveVI’s salient latent space (**Fig. 3d**). On the other hand, cell types separated clearly in contrastiveVI’s shared latent space (**Fig. 3e**). This result is consistent with prior knowledge, since we would expect the underlying factors of variation that distinguish cell types to be shared across healthy and infected cells. Moreover, we find both qualitatively (**Supplementary Fig. 3)** and quantitatively (**Fig. 3f**) that contrastiveVI’s shared latent space captures differences between cell types better than previously proposed contrastive methods’ shared latent spaces, with a similar degree of separation as the latent spaces of scVI and DCA. The disentanglement of contrastiveVI’s salient and shared latent factors was further corroborated by metrics previously proposed in the machine learning literature [30, 31, 32] for quantifying the degree of disentanglement between groups of latent factors (**Supplementary Fig. 4)**. Taken together, these results demonstrate that contrastiveVI successfully disentangles variations enriched in target data from shared variations, even when other methods struggle.

### Analyzing cell line responses to a small-molecule therapy

We next applied contrastiveVI to a dataset collected using the recently developed MIX-Seq [2] platform. MIX-Seq measures the transcriptional responses of up to hundreds of cancer cell lines in parallel after being treated with one or more small molecule compounds. Here our target dataset contains measurements collected by McFarland et al. [2] from 24 cell lines treated with idasanutlin. The small molecule idasanutlin is an antagonist of *MDM2*, a negative regulator of the tumor suppresor protein p53, hence offering cancer therapeutic opportunities [33]. Based on the mechanism of action of idasanutlin, activation of the p53 pathway is observed in cell lines with wild type *TP53* and not in transcriptionally inactive mutant *TP53* cell lines [33]. Our goal is thus to separate target cells by *TP53* mutation status. For the background dataset, we use measurements from the same cell lines treated with the control compound dimethyl sulfoxide (DMSO).

Qualitatively, contrastiveVI’s salient latent space stratifies cells based on *TP53* mutation status (**Fig. 4a**). In addition, our quantitative metrics indicate that contrastiveVI separates the two classes of target cells more clearly than baseline methods (**Fig. 4b**). We also used contrastiveVI to impute normalized expression levels of the p53-induced gene *TP53I3* for background cells as well as the two clusters of target cells identified by applying k-means clustering to the contrastiveVI salient latent space. We find clear differences in the imputed expression values for cells in the identified target cluster containing a majority of *TP53* wild type cells compared to background cells (**Fig. 4c**), which agrees with idasanutlin’s mechanism of action. In contrast, we find no discernible difference between imputed *TP53I3* expression values for background cells and the second cluster of target cells consisting almost exclusively of *TP53* mutant cells. To validate the quality of contrastiveVI’s imputations, we compared the values returned by contrastiveVI to those of scVI; we find that differences between scVI and contrastiveVI’s imputed expression values for both target and background cells are comparable to the differences in values between scVI models trained with different random initializations (**Supplementary Fig. 5)**.

**Figure 4:**
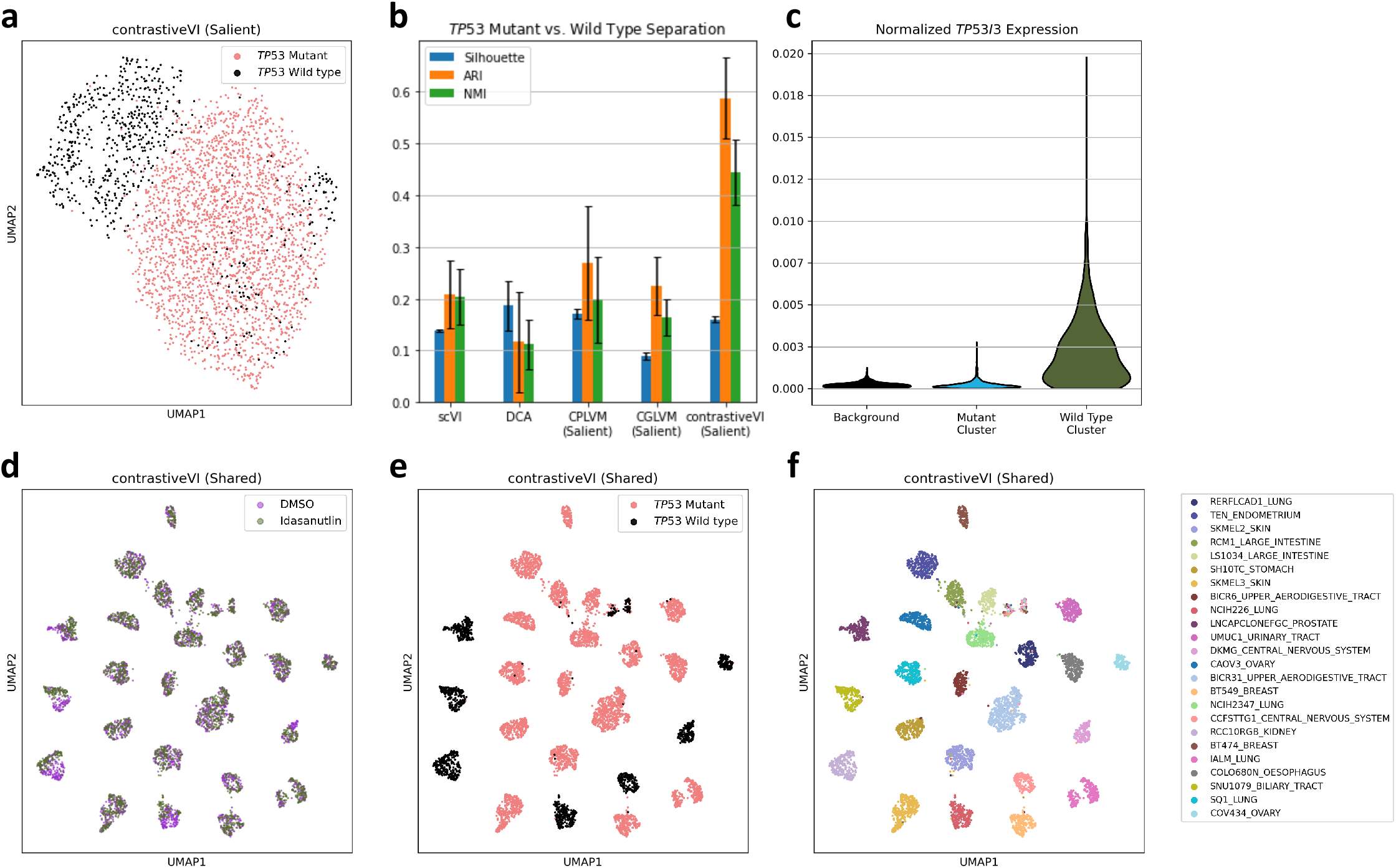
contrastiveVI stratifies cancer cells by response to idasanutlin. **a**, UMAP plot of contrastiveVI’s salient latent representations for idasanutlin-treated cells from McFarland et al. [2]. **b**, Clustering metrics quantify how well cells separate by *TP53* mutation status for scVI and DCA’s single latent spaces and contrastive models’ salient latent spaces, with means and standard errors across five random trials plotted. **c**, Normalized *TP53I3* expression levels returned by contrastiveVI for background cells and the two clusters of target cells identified in the contrastiveVI salient latent space. **d**,**e**,**f**, UMAP plots of contrastiveVI’s shared latent space colored by treatment type (**d**) *TP53* mutation status (**e**), and cell line (**f**).

We further explored the differences between the idasanutlin-treated cells compared to the controls via a modified version of the contrastiveVI differential expression test designed to compare a group of target cells with background cells (**Methods**). As done previously, we validated the results of this target-versus-background cell differential expression test by comparing the resulting Bayes factors with those returned by scVI models for the same task, and we find strong agreement between the two methods (**Supplementary Fig. 6)**. We find that the cluster containing a majority of *TP53* wild type cells has differentially expressed genes enriched in the p53 signaling pathway compared to background cells (**Supplementary Table 3**). In contrast, for the cluster associated with *TP53* mutation we find no enriched pathways when compared to background cells.

Finally, to understand the variations captured by contrastiveVI’s shared latent space, we embedded all cells, whether treated with DMSO or idasanutlin, into the model’s shared latent space. Ideally, contrastiveVI’s shared latent space would only capture variations that distinguish cell lines and not those related to treatment response. In particular, we would expect strong mixing between DMSO- and idasanutlin-treated cells even for cell lines with wild type *TP53*. We find that wild type *TP53* cell lines clearly separate by treatment type in the original data (**Supplementary Fig. 7)**, whereas cells mix more strongly across treatment types (**Fig. 4d**) regardless of *TP53* mutation status (**Fig. 4e**) and instead separate primarily by cell line (**Fig. 4f**) in the contrastiveVI shared latent space. These results further illustrate contrastiveVI’s ability to disentangle shared and target-specific variations.

### Isolating CRISPR-activation-induced variations in Perturb-Seq data

We next applied contrastiveVI to a Perturb-Seq [3, 34] dataset. Perturb-Seq combines high-throughput scRNA-seq methods with barcoding of CRISPR-induced genomic perturbations, enabling the evaluation of such perturbations at single-cell resolution. Previous studies have successfully leveraged Perturb-Seq to better understand regulatory circuits related to innate immunity [35], the unfolded protein response pathway [34], and the T cell receptor signaling pathway [36], among other applications. Despite these successes, recent work [7] suggested that naive approaches for analyzing Perturb-Seq data may fail to capture subtle perturbation-induced transcriptomic changes due to intercellular variations unrelated to the perturbations.

In this experiment we applied contrastiveVI to a Perturb-Seq dataset from Norman et al. [4]; the authors assessed the effects of 284 different CRISPR-mediated perturbations on the growth of K562 cells, where each perturbation induced the overexpression of a single gene or a pair of genes. Here, we focus on a subset of these perturbations for which the authors provided labels indicating a known gene program induced by the perturbation. We would expect cells to separate based on these gene programs; however, in the latent spaces of scVI and DCA models we observed significant mixing between cells with different gene program labels (**Fig. 5a-b**). We also observed significant mixing among gene programs in the salient latent spaces of CPLVM (**Fig. 5c**) and CGLVM (**Fig. 5d**) models trained using cells treated with control guides as a background dataset. In contrast, we find qualitatively (**Fig. 5e**) and quantitatively (**Fig. 5f**) that contrastiveVI better separates cells by gene program in its salient latent space. We note that our clustering metric values for this dataset are lower than those for previous datasets, potentially indicating that expression differences induced by the single- or double-gene CRISPR perturbations are more subtle than the clear separations found in previous datasets.

**Figure 5:**
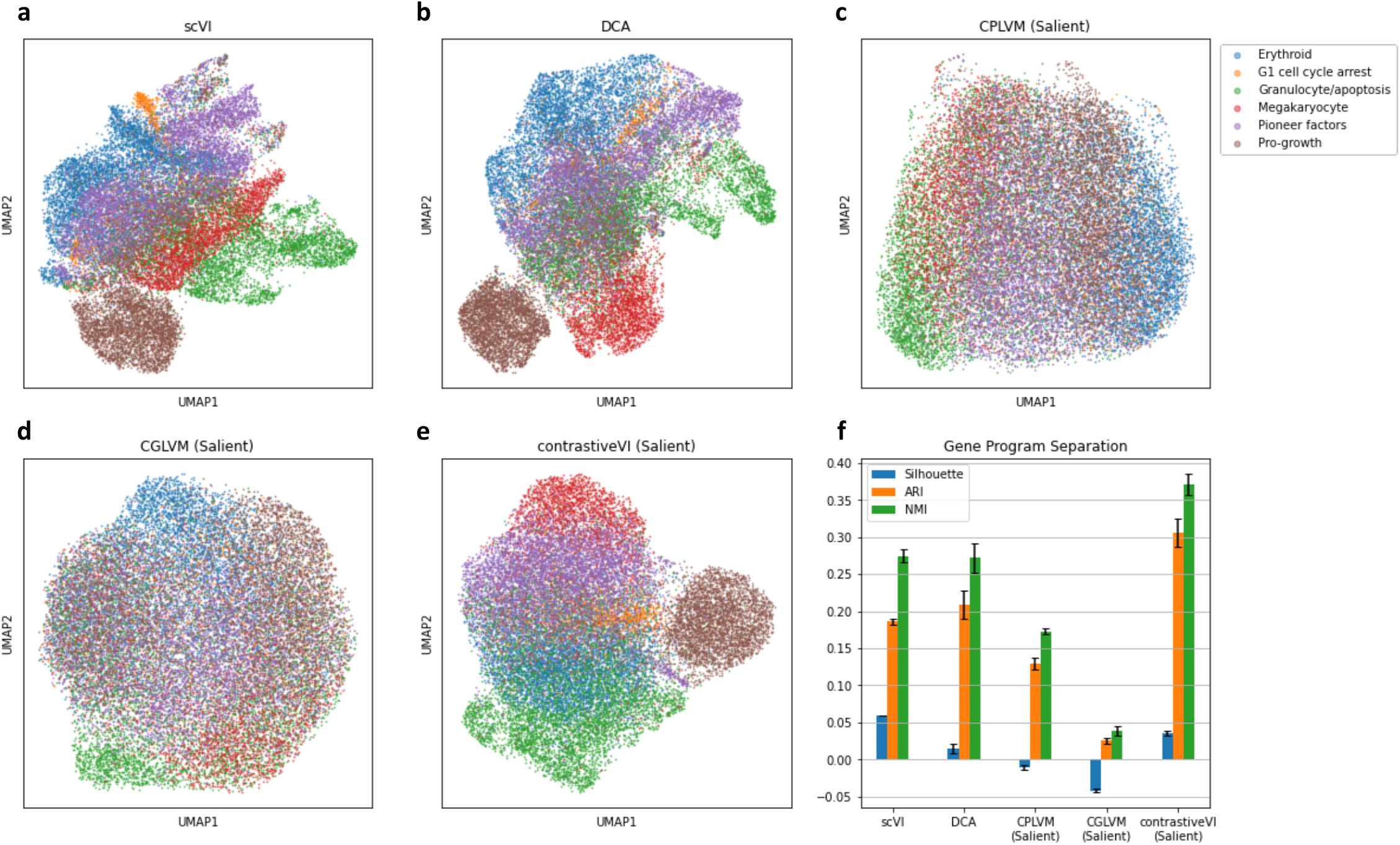
contrastiveVI isolates CRISPR-perturbation-induced variations in a large-scale Perturb-Seq experiment. **a-e**, UMAP plots colored by induced gene program of scVI’s latent space (**a**), DCA’s latent space (**b**), CPLVM’s salient latent space (**c**), CGLVM’s salient latent space (**d**), and contrastiveVI’s salient latent space (**e**). **f**, Clustering metrics quantifying how well cells separate by gene program label, with means and standard errors across five random trials reported for each method.

We also inspected contrastiveVI and baseline models’ shared latent spaces. Since the induced gene programs are specific to target (i.e., perturbed) cells, we would expect cells to mix by gene program in contrastive models’ shared latent spaces. We find qualitatively (**Supplementary Fig. 8a-e**) and quantitatively (**Supplementary Fig. 8f**) that contrastive models’ shared latent spaces exhibit strong mixing across gene programs as desired, with CGLVM and CPLVM slightly outperforming contrastiveVI quantitatively in this case.

### Extending contrastiveVI to multimodal single-cell datasets

ECCITE-Seq [24] is a recently developed platform that enables joint RNA and surface protein readouts of pooled CRISPR screens. Though previous works [23, 37, 38] have developed computational methods for integrating multi-omic measurements into unified representations of cellular state, they are not designed to isolate perturbation-induced variations in the data. Thus, these previously proposed models may learn representations of target cells dominated by irrelevant sources of variation shared with controls. To isolate the CRISPR-induced variations in ECCITE-Seq data, Papalexi et al. [20] proposed the mixscape algorithm. However, mixscape operates solely on transcriptomic data, and its results must be contextualized with protein information post hoc. This approach is not ideal, since it biases any analyses towards transcriptome measurements, which may not be sufficient for differentiating functionally diverse cell populations [23, 38].

To address this gap in methodology, we extended totalVI, a deep generative model designed for the analysis of paired RNA and protein measurements, using the contrastive latent variable modeling techniques employed in contrastiveVI. We then applied our resulting totalContrastiveVI model (**Methods**) to analyze an ECCITE-Seq dataset collected in Papalexi et al. [20]. In that work, the authors sought to explore the regulatory networks underlying the expression of immune checkpoint molecules, such as programmed death-ligand 1 (PD-L1), in THP-1 [39] cells. To do so, they measured cells’ transcriptomes alongside surface protein levels of the proteins PD-L1, PD-L2, CD86 and CD366 for cells perturbed via one of 111 CRISPR guides as well as for a set of control cells. As a baseline, we first applied totalVI [23] to learn a lower-dimensional representation of the perturbed cells. Ideally the model would capture perturbation-induced variations; however, we find instead that variations in the totalVI latent space are dominated by nuisance factors, such as transduction replicate identity and cell cycle stage (**Fig. 6a**, top row). Using measurements from the control cells as a background, we next applied totalContrastiveVI to this dataset. As expected, we find that the totalContrastiveVI shared latent space is also dominated by nuisance variations (**Supplementary Fig. 9a-b**). In contrast, the totalContrastiveVI salient latent space exhibits a clear clustering structure invariant to replicate identity and cell cycle stage (**Fig. 6a**, bottom row). Of the three clusters revealed in the salient latent space of totalContrastiveVI, we find that one consists of cells perturbed for upstream components of the IFN-*γ* pathway, one consists solely of cells perturbed for *IRF1*, which encodes for an IFN-*γ* mediator, and the remaining cluster consists of cells from other perturbations (**Fig. 6b**).

**Figure 6:**
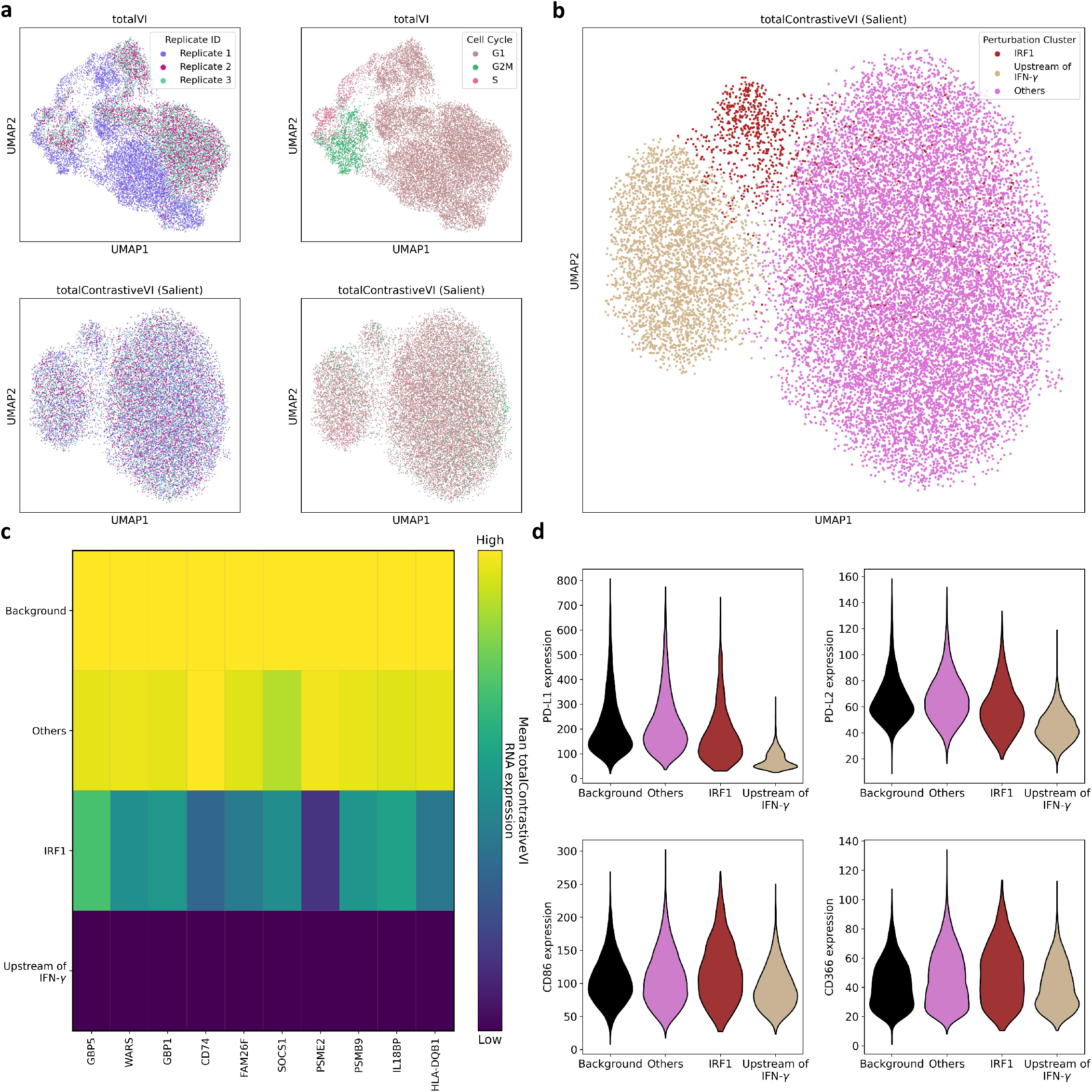
totalContrastiveVI isolates perturbation-induced variations in joint RNA and protein measurements. **a**, totalVI embeddings (top) and totalContrastiveVI salient embeddings (bottom) colored by replicate number and cell cycle stage. **b**, The three clusters revealed in the totalContrastiveVI salient latent space colored by perturbation category. **c**, Mean imputed normalized RNA expression levels of IFN-*γ* pathway genes for background cells and the three clusters of perturbed cells identified in the totalContrastiveVI salient latent space. **d**, Imputed protein expression levels for background cells and the three clusters of perturbed cells.

We then used totalContrastiveVI to better understand the differences between these three groups of target cells and the unperturbed background cells. We began by applying totalContrastiveVI to impute normalized RNA values for genes in the IFN-*γ* pathway (**Fig. 6c**). We find that these genes are downregulated in cells perturbed for upstream components of the IFN-*γ* pathway compared to unperturbed cells and those perturbed for genes unrelated to the IFN-*γ* pathway. These genes also appear to be downregulated in cells perturbed for *IRF*1, though not as strongly as for cells perturbed for upstream components of the IFN-*γ* pathway. Furthermore, we imputed denoised protein expression values for the four proteins measured in this dataset (**Fig. 6d**) using totalContrastiveVI. Cells perturbed for upstream components of the IFN-*γ* pathway seem to exhibit a marked decrease in PD-L1 protein levels compared to unperturbed cells and cells perturbed for genes unrelated to IFN-*γ*, and we verified these differences using totalContrastiveVI to perform a protein differential expression test (**Methods**). We also found a smaller but noticeable decrease in PD-L2 protein levels for cells perturbed for upstream components of the IFN-*γ* pathway. In contrast, these perturbations did not have a notable effect on cells’ CD86 and CD366 protein levels. In addition, despite the changes in RNA levels found for cells perturbed for *IRF*1, these cells do not exhibit noticeable changes in protein expression levels compared to unperturbed cells.

These results demonstrate that, beyond unimodal scRNA-seq datasets, contrastive latent variable modeling can facilitate better understanding of variations enriched in multimodal target single-cell datasets.

## Discussion

In this work we introduced contrastiveVI, a framework for explicitly disentangling enriched variations in a target single-cell dataset from those shared with a related background dataset using deep contrastive latent variable models. contrastiveVI is the first method designed to analyze single-cell count data in the contrastive analysis setting that both directly models the technical factors of variation in single-cell data and leverages the expressive power of deep learning. Moreover, to demonstrate how our framework can be easily extended to multimodal single-cell datasets, we also developed totalContrastiveVI for isolating enriched variations in joint RNA and surface protein measurements. Our models provide many capabilities out of the box, such as dimensionality reduction, imputation, and differential expression testing, to enhance our understanding of the biological processes underlying the salient variations in a given target dataset.

In four different contexts—response to cancer treatment, infection by different pathogens, exposure to drug compounds, and genomic perturbation via CRISPR guides—we find that contrastiveVI successfully isolates enriched variations in target cells’ transcriptomes. Similarly, we find that our multimodal totalContrastiveVI model successfully isolates enriched variations in a joint RNA and surface protein dataset. Given the continuing development of new sequencing technologies for efficiently measuring cellular responses to many perturbations in parallel, we expect our models to be of immediate interest to the single-cell research community. The models are available as an open-source software package and are implemented using the scvi-tools [40] Python library, thereby enabling seamless interoperability with the Scanpy [41] and Seurat [15] software ecosystems.

The ideas underpinning contrastiveVI admit multiple potential directions for future work. Our contrastive modeling approach could be further extended to handle additional modalities beyond RNA transcript and protein counts by integrating previously proposed modality-specific likelihoods, such as that of peakVI [42] for chromatin accessibility, into the contrastiveVI framework. The incorporation of further modalities could provide a more complete picture of changes in cellular state due to various perturbations. Moreover, recent work [43, 44] has demonstrated success in learning interpretable representations of single-cell data where each dimension of the representation corresponds to a known gene pathway. Incorporating such constraints into contrastiveVI could shed further light on the biological phenomena captured in the salient and shared latent spaces learned by complex contrastiveVI models.

## Methods

### The contrastiveVI model

Here, we present the contrastiveVI model in more detail. We begin by describing the model’s generative process and then the model’s inference procedure.

### The contrastiveVI generative process

For a target data point *x*_*n*_, we assume that each expression value *x*_*ng*_ for sample *n* and gene *g* is generated through the following process:

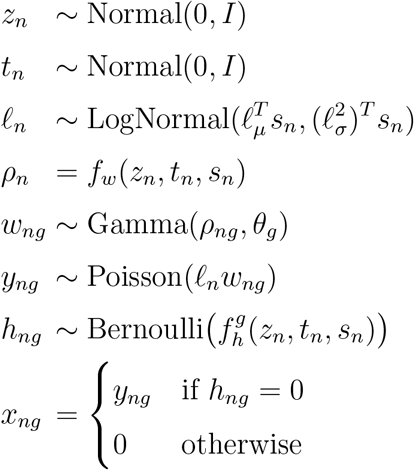

In this process, *z*_*n*_ and *t*_*n*_ refer to the two sets of latent variables underlying variations in scRNA-seq expression data. Here, *z*_*n*_ represents variables that are shared across background and target cells, while *t*_*n*_ represents variations unique to target cells. We place a standard multivariate Gaussian prior on both sets of latent factors, since this specification is computationally convenient for inference in the VAE framework [25]. To encourage the disentanglement of latent factors, for background data points *b*_*n*_, we assume the same generative process but instead set *t*_*n*_ = **0** to represent the absence of salient latent factors in the generative process. Categorical covariates, such as experimental batches, are represented by *s*_*n*_.

Here, 𝓁_*μ*_ and 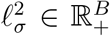, where *B* denotes the cardinality of the categorical covariate, parameterize the prior for latent RNA library size scaling factor on a log scale, and *s*_*n*_ is a *B*-dimensional one-hot vector encoding a categorical covariate index. For each category (e.g., experimental batch), 𝓁_*μ*_ and 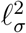 are set to the empirical mean and variance of the log library size, respectively. The gamma distribution is parameterized by the mean *ρ*_*ng*_ ∈ ℝ _+_ and shape *θ*_*g*_ ∈ ℝ _+_. Furthermore, following the generative process, *θ*_*g*_ is equivalent to a gene-specific inverse dispersion parameter for a negative binomial distribution, and 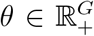 is estimated via variational Bayesian inference. *f*_*w*_ and *f*_*g*_ in the generative process are neural networks that transform the latent space and batch annotations to the original gene space, i.e.: ℝ^*d*^ × {0, 1}^*B*^ → ℝ^*G*^, where *d* is the size of the concatenated salient and shared latent spaces. The network *f*_*w*_ is constrained during inference to encode the mean proportion of transcripts expressed across all genes using a softmax activation function in the last layer. That is, letting 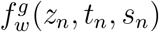 denote the entry in the output of *f*_*w*_ corresponding to gene *g*, we have 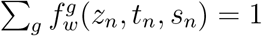. The neural network *f*_*h*_ encodes whether a particular gene’s expression has dropped out in a cell due to technical factors.

Our generative process closely follows that of scVI [11], with the addition of the salient latent factors *t*_*n*_. While scVI’s modeling approach has been shown to excel at many scRNA-seq analysis tasks, our empirical results demonstrate that it is not suited for contrastive analysis (CA). By dividing the RNA latent factors into shared factors *z*_*n*_ and target-specific factors *t*_*n*_, contrastiveVI successfully isolates variations enriched in target datasets that were missed by previous methods. We depict the full contrastiveVI generative process as a graphical model in **Supplementary Fig. 10**.

### Inference with contrastiveVI

We cannot compute the contrastiveVI posterior distribution using Bayes’ rule because the integrals required to compute the model evidence *p*(*x*_*n*_|*s*_*n*_) are analytically intractable. As such, we instead approximate our posterior distribution using variational inference [45]. For target data points, we approximate our posterior with a distribution factorized as follows:

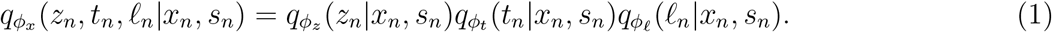

Here, *ϕ*_*x*_ denotes a set of learned weights used to infer the parameters of our approximate posterior. Based on our factorization, we can divide *ϕ*_*x*_ into three disjoint sets *ϕ*_*z*_, *ϕ*_*t*_ and *ϕ*_𝓁_ for inferring the parameters of the distributions of *z, t* and 𝓁, respectively. Following the VAE framework [25], we then approximate the posterior for each factor as a deep neural network that takes as input expression levels and outputs the parameters of its corresponding approximate posterior distribution (e.g., mean and variance). Moreover, we note that each factor in the posterior approximation shares the same family as its respective prior distribution (e.g., *q*(*z*_*n*_|*x*_*n*_, *s*_*n*_) follows a normal distribution). We can simplify our likelihood by integrating out *w*_*ng*_, *h*_*ng*_, and *y*_*ng*_, yielding *p*_*ν*_(*x*_*ng*_|*z*_*n*_, *t*_*n*_, *s*_*n*_, 𝓁_*n*_), which follows a zero-inflated negative binomial (ZINB) distribution (**Supplementary Note 3**) and where *ν* denotes the parameters of our generative model. As with our approximate posteriors, we realize our generative model with deep neural networks. For Equation 1 we can derive (**Supplementary Note 4**) a corresponding variational lower bound:

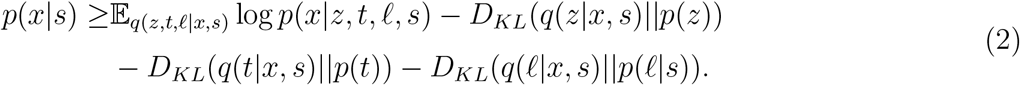

Next, for background data points we approximate the posterior using the factorization:

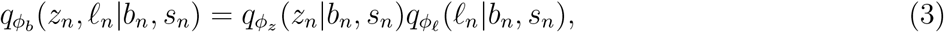

where *ϕ*_*b*_ denotes a set of learned parameters used to infer the values of *z*_*n*_ and 𝓁_*n*_ for background samples. Following our factorization, we divide *ϕ*_*b*_ into the disjoint sets *ϕ*_*z*_ and *ϕ*_𝓁_. We note that *ϕ*_*z*_ and *ϕ* are shared across target and background samples; this encourages the posterior distributions 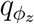 and 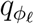 to capture variations shared across the target and background cells, while 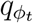 captures variations unique to the target data. Once again, we can simplify our likelihood by integrating out *w*_*ng*_, *h*_*ng*_, and *y*_*ng*_ to obtain *p*_*ν*_(*x*_*ng*_|*z*_*n*_, **0**, *s*_*n*_, 𝓁_*n*_), which follows a ZINB distribution. We similarly note that the parameters of our generative model *ν* are shared across target and background points to encourage *z* to capture shared variations across target and background points while *t* captures target-specific variations. We then have the following variational lower bound for our background data points:

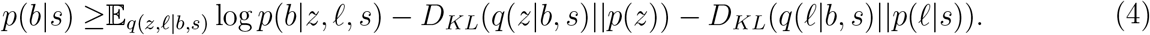

We then jointly optimize the parameters of our generative model and inference networks using stochastic gradient descent to maximize the sum of these two bounds over our background and target data points. All neural networks used to implement the variational and generative distributions were feedforward and used standard activation functions. We used the same network architecture and hyperparameter values for all experiments, and we refer the reader to **Supplementary Note 5** for more details.

### Differential gene expression analysis with contrastiveVI

For two cell groups *A* = (*a*_1_, *a*_2_, …, *a*_*n*_) and *B* = (*b*_1_, *b*_2_, …, *b*_*m*_) in the target dataset, the posterior probability of gene *g* being differentially expressed in the two groups can be obtained as proposed by Boyeau et al. [46]. For any arbitrary cell pair *a*_*i*_, *b*_*j*_, we have two mutually exclusive models:

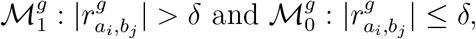

where 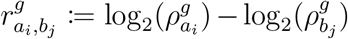 is the log fold change of the denoised, library-size-normalized expression of gene *g*, and *δ* is a pre-defined threshold for log fold change magnitude to be considered biologically meaningful. The posterior probability of differential expression is therefore expressed as 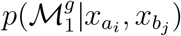, which can be obtained via marginalization of the latent variables and categorical covariates:

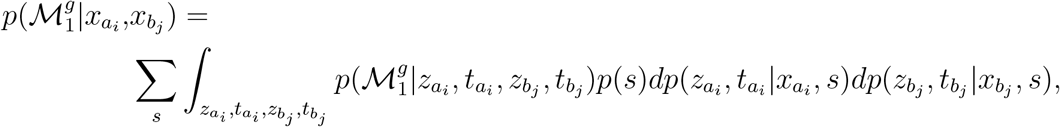

where *p*(*s*) is the relative abundance of target cells in category *s*, and the integral can be computed via Monte Carlo sampling using the variational posteriors 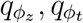. Finally, the group-level posterior probability of differential expression is

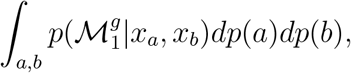

where we assume that the cells *a* and *b* are independently sampled *a* ∼ *𝒰* (*a*_1_, …, *a*_*m*_) and *b* ∼ *𝒰* (*b*_1_, …, *b*_*m*_), respectively. Computationally, this quantity can be estimated by a large random sample of pairs from the cell group *A* and *B*.

This procedure, with a minor modification, can also be used to test for differentially expressed genes between a group of target cells and a group of background cells. Without loss of generality, let *A* denote a group of cells in the target dataset and *B* denote a group of cells in the background dataset. When computing the integral in the expression for 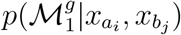, the values of 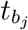 are fixed at **0** to represent their absence in the generative process for background cells. The test then proceeds as previously described for the case of two groups of target cells.

In our experiments, 10,000 cell pairs were sampled, 100 Monte Carlo samples were obtained from the variational posteriors for each cell, and the *δ* threshold was set to 0.25, the default value recommended by the scvi-tools Python library [40]. Genes with a group-level posterior probability of differential expression greater than 0.95 were considered for downstream pathway enrichment analysis.

### The totalContrastiveVI model

We now present the totalContrastiveVI model in more detail.

#### The totalContrastiveVI generative process

Formally, for a given cell *n*, we have gene expression values *x*_*ng*_ for each measured gene *g* and protein expression values *y*_*nτ*_ for each measured protein *τ*. For gene expression values, we assume the generative process described previously for contrastiveVI.

To account for the technical biases of CITE-Seq-based platforms, totalContrastiveVI models protein counts as a mixture of foreground and background components. For target cells, the full generative process for protein measurements is as follows:

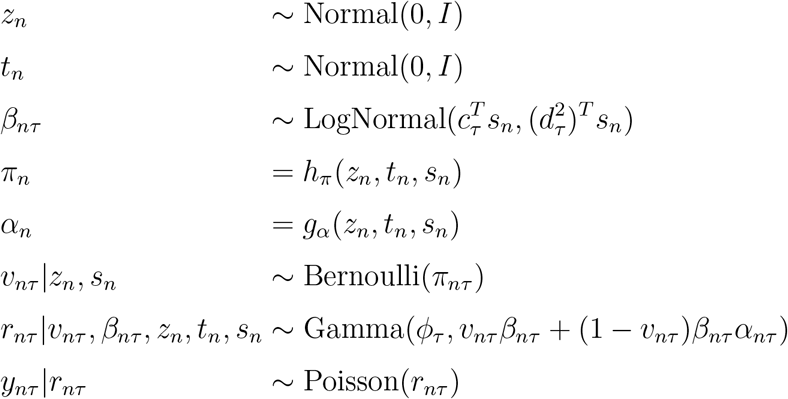

Here, *β*_*nτ*_ is a protein-specific variable representing a protein-specific background intensity. The parameters *c*_*τ*_ ∈ ℝ^*B*^ and 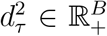 for the prior distribution of *β*_*nτ*_ are protein-specific and treated as model parameters to be learned during training. *ν*_*nτ*_ controls whether a given protein’s counts are generated from the background or foreground mixture component, with its parameter *π*_*nτ*_ being the output of the neural network *h*_*π*_ and representing the probability of the counts being generated due to background alone. *g*_*α*_ is constrained such that its output *α*_*nτ*_ always exceeds 1. Thus, one of the mixture components will always be larger than the other, enabling one to be interpreted as foreground and the other as background. For a given mixture component, *y*_*nτ*_ |*z*_*n*_, *t*_*n*_, *s*_*n*_, *β*_*nτ*_ follows a negative binomial distribution, which can be shown by integrating out *r*_*nτ*_. Moreover, *y*_*nτ*_ given *z*_*n*_, *t*_*n*_, and *s*_*n*_ can be shown to follow a negative binomial distribution by integrating out *v*_*nτ*_, with *ϕ*_*τ*_ acting as a protein-specific inverse dispersion parameter. For background data points, we assume the same generative process but set *t*_*n*_ = **0** to represent the absence of salient latent factors.

The generative process of totalContrastiveVI closely follows that of totalVI [23], but with the addition of salient latent factors. We depict the totalContrastiveVI generative process as a graphical model in **Supplementary Fig. 11**.

#### Inference with totalContrastiveVI

As with contrastiveVI, for totalContrastiveVI we approximate our posterior distribution using variational inference. For target data points we use an approximate posterior factorized as follows:

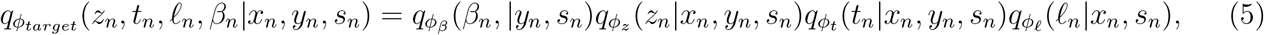

where *ϕ*_*target*_ denotes a set of learned weights for our approximate posterior distribution. We can simplify the gene and protein likelihood as described previously to obtain *p*_*v*_(*x*_*ng*_, |*z*_*n*_, *t*_*n*_, 𝓁_*n*_, *s*_*n*_), which is a zero-inflated negative binomial distribution, and *p*_*v*_(*y*_*nt*_, |*z*_*n*_, *t*_*n*_, *β*_*n*_, *s*_*n*_), which is a mixture of negative binomials.

For background points we have the following approximate posterior distribution:

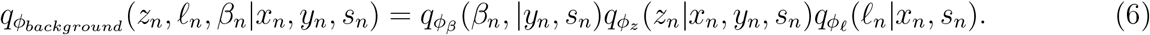

We then jointly optimize the parameters of our generative model and inference networks using stochastic gradient descent to maximize the sum of the ELBOs over our background and target data points. All neural networks used to implement the variational and generative distributions were feedforward and used standard activation functions. We used the same network architecture and hyperparameter values for all experiments, and we refer the reader to **Supplementary Note 6** for more details.

#### Differential expression analysis with totalContrastiveVI

For two groups of cells *A* = (*a*_1_, *a*_2_, …, *a*_*n*_) and *B* = (*b*_1_, *b*_2_, …, *b*_*m*_) in the target dataset, totalContrastiveVI can be used to compute both differentially expressed genes and differentially expressed protiens. To compute differentially expressed genes, we use the same procedure as in contrastiveVI. For proteins, we use the same framework, with the following two mutually exclusive hypotheses given two cells *a*_*i*_ and *b*_*j*_:

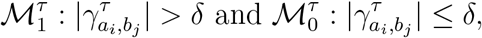

where

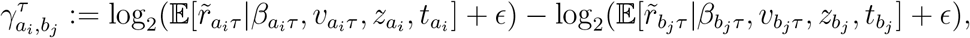

and our conditional expectation is

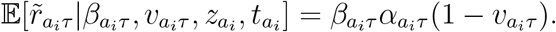

Based on our previous definition of *v*, this expression can be interpreted as the foreground mean if the cell was generated from the foreground, and zero otherwise. The value *ϵ* serves as a “prior count” that ensures the log fold change is defined even when the conditional expectation is zero. In our experiments we set *ϵ* = 0.5, as was done in Gayoso et al. [23]. Let *C*_*n*_ = {*x*_*n*_, *y*_*n*_, *s*_*n*_} denote the observed data for a cell *n*. The posterior probability of differential expression is therefore expressed as 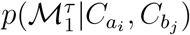, which can be obtained by integrating with respect to the distribution

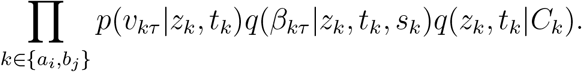

The group-level posterior probability of differential expression can then be estimated by sampling, as previously described for the contrastiveVI gene differential expression test. For our protein differential expression tests, 10,000 cell pairs were sampled, 100 Monte Carlo samples were obtained from the variational posteriors for each cell, and the *δ* threshold was set to 0.15.

### Pathway enrichment analysis

Pathway enrichment analysis refers to a computational procedure for determining whether a predefined set of genes (i.e., a gene pathway) has statistically significant differences in expression between two biological states. Many tools exist for performing pathway enrichment analysis (see [47] for a review). Our analyses used Enrichr [48], a pathway analysis tool for non-ranked gene lists based on Fisher’s exact test, to find enriched pathways from the KEGG pathway database [28]. Specifically, the Enrichr wrapper implemented in the open-source GSEAPy^1^ Python library was used for our analyses. Pathways enriched at false discovery rate smaller than 0.05—adjusted by the Benjamini-Hochberg procedure [49]— are reported in this study.

### Baseline models

As discussed in the main text, due to previous findings indicating that normalization of scRNA-seq data has a substantial impact on downstream results [12], we only consider CA methods specifically tailored for scRNA-seq count data as baselines in this study. To our knowledge, CPLVM (contrastive Poisson latent variable model) and CGLVM (contrastive generalized latent variable model) are the only CA methods that explicitly model count-based scRNA-seq normalization [7]. We present a summary of previous work in CA in **Supplementary Table 4**. To illustrate the need for models specifically designed for CA, we also consider scVI, a deep generative model for scRNA-seq count data that takes batch effect, technical dropout, and varying library size into modeling consideration [11], as well as deep count autoencoder (DCA), an autoencoder neural network for reducing noise in scRNA-seq count data due to technical dropout [27]. Below, we describe the CA methods CPLVM and CGLVM in more detail.

In CPLVM, variations shared between the background and target conditions are assumed to be captured by the shared latent variable values 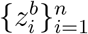 and 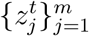, and target-condition-specific variations are captured by the salient latent variable values 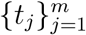, where *n, m* are the number of background and target cells, respectively. Library size differences between the two conditions are modeled by 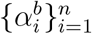 and 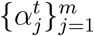, whereas gene-specific library sizes are parameterized by 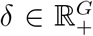, where *G* is the number of genes. Each data point is considered Poisson distributed, with the rate parameter determined by 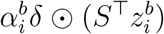 for a background cell *i* and by 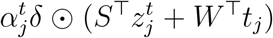 for a target cell *j*, where *S, W* are model weights that linearly combine the latent variables, and ⨀ represents an element-wise product. The model weights and latent variables are assumed to have Gamma priors, *δ* has a standard log-normal prior, and 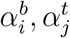 have log-normal priors with parameters given by the empirical mean and variance of log total counts in each dataset. Posterior distributions are fitted using variational inference with mean-field approximation and log-normal variational distributions.

The CA modeling approaches of CGLVM and CPLVM are similar. In CGLVM, however, the relationships of latent factors are considered additive and relate to the Poisson rate parameter via an exponential link function (similar to a generalized linear modeling scheme). All the priors and variational distributions are Gaussian in CGLVM.

### Model optimization details

For all datasets, contrastiveVI or totalContrastiveVI models were trained with 80% of the background and target data; the remaining 20% was reserved as a validation set for early stopping to determine the number of training epochs needed. Training was early stopped when the validation variational lower bound showed no improvement for 45 epochs, typically resulting in 127 to 500 epochs of training. All contrastiveVI and totalContrastiveVI models were trained with the Adam optimizer [50], with *ε* = 0.01, learning rate at 0.001, and weight decay at 10^−6^. The same hyperparameters and training scheme were used to optimize the scVI models using only target data, usually with 274 to 500 epochs of training based on the early stopping criterion. As in the open-source implementation by Eraslan et al., DCA models were trained for a maximum of 500 epochs using the RMSprop optimizer with a learning rate at 0.001, with early stopping when the validation loss showed no improvement for 15 epochs [27]. As in Jones et al., the CPLVMs were trained via variational inference using all background and target data for 2,000 epochs with the Adam optimizer with *ε* = 10^−8^ and a learning rate at 0.05, and the CGLVMs were similarly trained for 1,000 epochs with a learning rate at 0.01 [7]. All models were trained with 10 salient and 10 shared latent variables five times with different random weight initializations. To understand the impact of the size of the salient latent space on model performance, we also trained models with varying salient latent dimension sizes and obtained overall consistent results (**Supplementary Fig. 12)**.

### Datasets and preprocessing

We now briefly describe all datasets used in this work along with any corresponding preprocessing steps. For our experiments datasets were chosen that not only had cells in a target and corresponding background condition, but also which had ground truth subclasses of target cells. Moreover, to avoid potential confounding effects, datasets collected using a variety of single-cell platforms (**Supplementary Table 5**) were used in our experiments. All preprocessing steps were performed using the Scanpy Python package [41], and all of our code for downloading and preprocessing these datasets is publicly available at https://github.com/suinleelab/contrastiveVI-reproducibility. For all experiments we retained the top 2,000 most highly variable genes returned from the Scanpy `highly_variable_genes` function, with the `flavor` parameter set to `seurat_v3`. For all datasets, the number of cells in the background vs. target condition can be found in **Supplementary Table 5**.

#### Zheng et al., 2017

This dataset consists of single-cell RNA expression levels of a mixture of bone marrow mononuclear cells (BMMCs) from 10x Genomics.^2^ For our target dataset, we use samples taken from patients with acute myeloid leukemia (AML) before and after a stem cell transplant. For our background dataset, we use measurements taken from two healthy control patients released as part of the same study. All data is publicly available: files containing measurements from the first patient pre- and post-transplant can be found https://cf.10xgenomics.com/samples/cell-exp/1.1.0/aml027_pre_transplant/aml027_pre_transplant_filtered_gene_bc_matrices.tar.gz and https://cf.10xgenomics.com/samples/cell-exp/1.1.0/aml027_post_transplant/aml027_post_transplant_filtered_gene_bc_matrices.tar.gz, respectively; from the second patient pre- and post-transplant https://cf.10xgenomics.com/samples/cell-exp/1.1.0/aml035_pre_transplant/aml035_pre_transplant_filtered_gene_bc_matrices.tar.gz and https://cf.10xgenomics.com/samples/cell-exp/1.1.0/aml035_post_transplant/aml035_post_transplant_filtered_gene_bc_matrices.tar.gz, respectively; and from the two healthy control patients https://cf.10xgenomics.com/samples/cell-exp/1.1.0/frozen_bmmc_healthy_donor1/frozen_bmmc_healthy_donor1_filtered_gene_bc_matrices.tar.gz and https://cf.10xgenomics.com/samples/cell-exp/1.1.0/frozen_bmmc_healthy_donor2/frozen_bmmc_healthy_donor2_filtered_gene_bc_matrices.tar.gz.

#### Haber et al., 2017

This dataset (Gene Expression Omnibus accession number GSE92332) used scRNA-seq measurements to investigate the responses of intestinal epithelial cells in mice to different pathogens. Specifically, in this dataset, responses to the bacterium *Salmonella* and the parasite *H. polygyrus* were investigated. Our target dataset included measurements of cells infected with *Salmonella* and from cells 10 days after being infected with *H. polygyrus*, while our background consisted of measurements from healthy control cells released as part of the same study. The number of cells of each cell type can be found in **Supplementary Table 6**.

#### McFarland et al., 2020

This dataset measured cancer cell lines’ transcriptional responses after being treated with various small-molecule therapies. For our target dataset, we used data from cells that were exposed to idasanutlin, and for our background we used data from cells that were exposed to a control solution of dimethyl sulfoxide (DMSO). *TP53* mutation status was determined using the DepMap [51] 19Q3 data release, available at https://depmap.org/. The count data was downloaded from the authors’ Figshare repository at https://figshare.com/articles/dataset/MIX-seq_data/10298696. The number of cells for each cell line can be found in **Supplementary Table 7**.

#### Norman et al., 2019

This dataset (Gene Expression Omnibus accession number GSE133344) measured the effects of 284 different CRISPR-mediated perturbations on K562 cells, where each perturbation induced the over-expression of a single gene or a pair of genes. As done in the analysis from Norman et al. [4], we excluded cells with the perturbation label `NegCtrl1_NegCtrl0__NegCtrl1_NegCtrl0` from our analysis. For our background dataset, we used all remaining unperturbed cells; for our target dataset, we used all perturbed cells that had a gene program label provided by the authors.

#### Papalexi et al., 2021

This dataset (Gene Expression Omnibus accession number GSE153056) measured the effects of 111 different CRISPR knockout perturbations on THP-1 cells. The dataset contains both transcriptomic measurements and measurements of surface protein levels for the proteins CD86, PD-L1, PD-L2, and CD366. Our background dataset consists of measurements from cells infected with non-targeting (NT) guide RNAs, while our target dataset consists of measurements from the perturbed cells.

### Evaluation Metrics

Here, we describe the quantitative metrics used in this study. All metrics were computed using their corresponding implementations in the scikit-learn Python package [52].

#### Silhouette width

We calculate silhouette width using the latent representations returned by each method. For a given sample *i*, the sillhouete width *s*(*i*) is defined as follows. Let *a*(*i*) be the average distance between *i* and other samples with the same ground truth label, and let *b*(*i*) be the smallest average distance between *i* and all other samples with a different label. The silhouette score *s*(*i*) is then

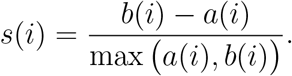

A silhouette width close to one indicates that *i* is tightly clustered with cells having the same ground truth label, while a score close to -1 indicates that a cell has been grouped with cells having a different label.

#### Adjusted Rand index

The adjusted Rand index (ARI) measures agreement between reference clustering labels and labels assigned by a clustering algorithm. Given a set of *n* samples and two sets of clustering labels describing those cells, the overlap between clustering labels can be described using a contingency table, where each entry indicates the number of cells in common between the two sets of labels. Mathematically, the ARI is calculated as

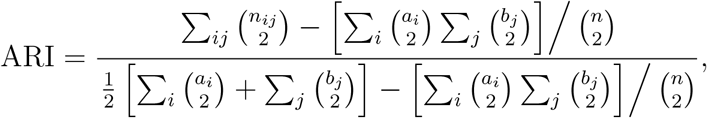

where *n*_*ij*_ is the number of cells assigned to cluster *i* based on the reference labels and cluster *j* based on a clustering algorithm, *a*_*i*_ is the number of cells assigned to cluster *i* in the reference set, and *b*_*j*_ is the number of cells assigned to cluster *j* by the clustering algorithm. ARI values closer to 1 indicate stronger agreement between the reference labels and labels assigned by a clustering algorithm.

#### Normalized mutual information

The normalized mutual information (NMI) measures the agreement between reference clustering labels and labels assigned by a clustering algorithm. The NMI is calculated as

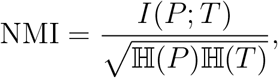

where *P* and *T* denote empirical distributions for the predicted and true clusterings, *I* denotes mutual information, and ℍ the Shannon entropy.

## Code Availability

Our Python software package with implementations of the contrastiveVI and totalContrastiveVI models is available at https://github.com/suinleelab/contrastiveVI. Code for reproducing the specific results in this paper is available at https://github.com/suinleelab/contrastiveVI-reproducibility.

## Data Availability

All datasets analyzed in this paper are publicly available. Our code for downloading and preprocessing them is available at https://github.com/suinleelab/contrastiveVI-reproducibility.

## Footnotes

1 https://gseapy.readthedocs.io/en/latest/

2 https://support.10xgenomics.com/single-cell-gene-expression/datasets

