## Supplementary materials for "Isolating salient variations of interest in single-cell data with contrastiveVI"

### Supplementary Figures

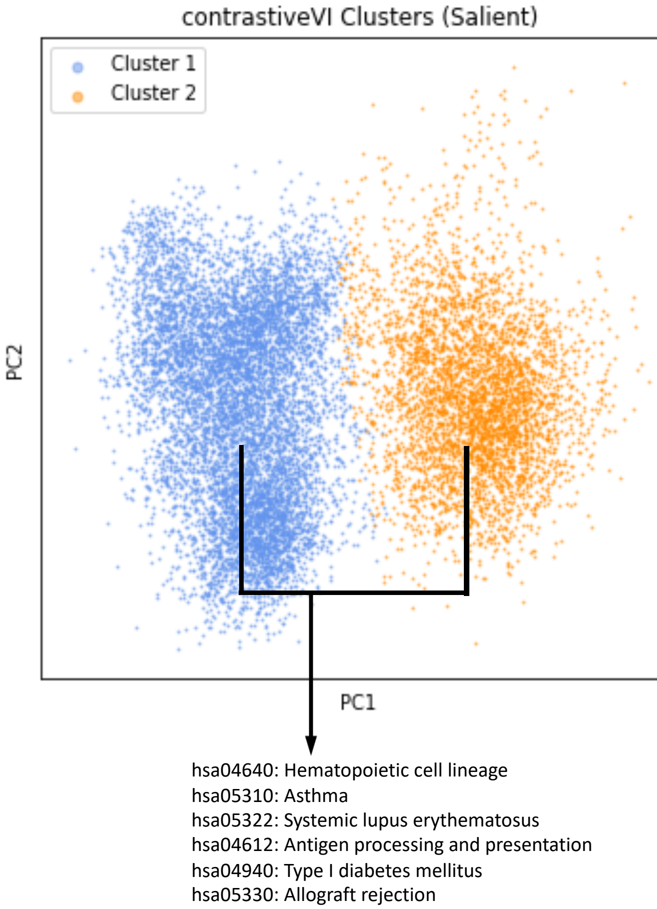

**Supplementary Figure 1: Clustering and pathway enrichment analysis using the salient latent space of contrastiveVI for BMMCs (bone marrow mononuclear cells) from AML (acute myeliod leukemia) patients.** contrastiveVI's salient latent representations of the Zheng et al. [19] target dataset (BMMCs from two AML patients before and after an allogenic stem-cell transplant) were clustered into two groups, and pathway enrichment analysis was then performed on the differentially expressed genes between the two clusters.

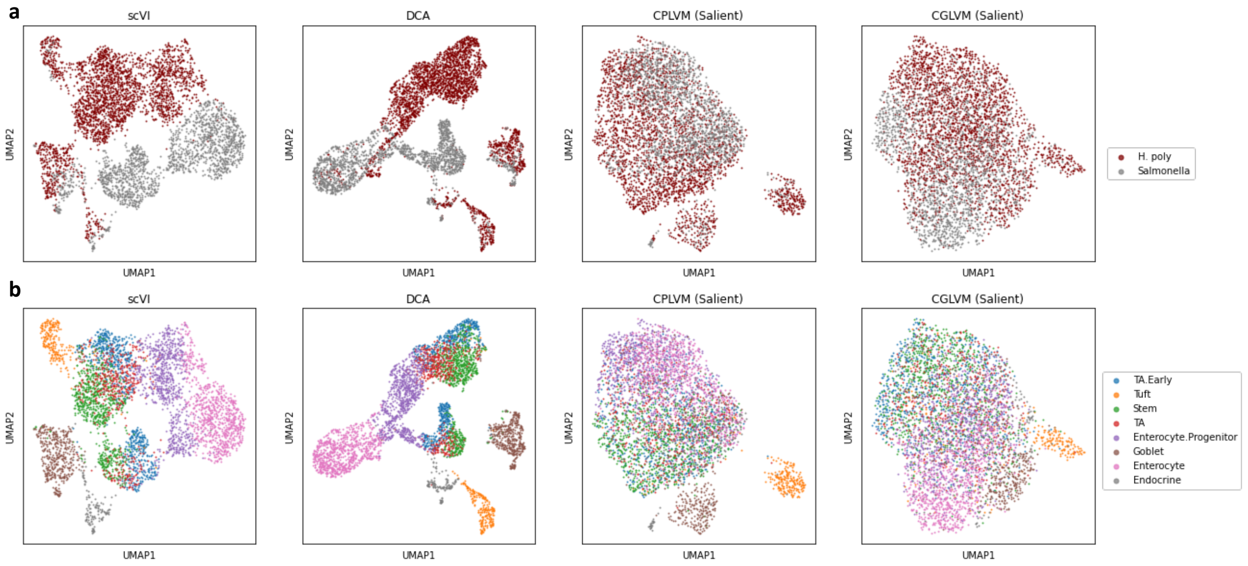

**Supplementary Figure 2: Salient latent spaces for mouse intestine epithelial cells. a,b,** UMAP plots of the salient latent spaces for the mouse intestine epithelial cell dataset colored by infection type (**a**) and cell type (**b**). For scVI and DCA, a UMAP plot of the model's single latent space is provided. We find that CPLVM and CGLVM do not separate cells by infection type as well as contrastiveVI. While scVI and DCA do have some separation by infection type, they strongly entangle the infection-induced variations with those due to cell type differences.

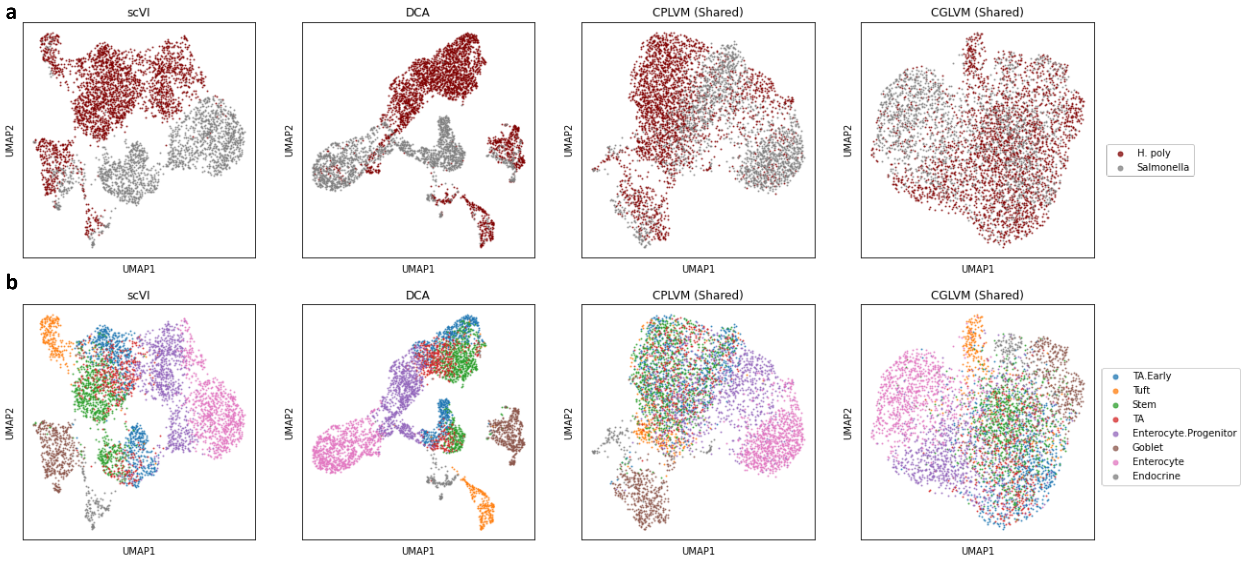

**Supplementary Figure 3: Shared latent spaces for mouse intestine epithelial cells.** **a,b,** UMAP plots of the shared latent spaces for the mouse intestine epithelial cell dataset colored by infection type (**a**) and cell type (**b**). For scVI and DCA, a UMAP plot of the model's single latent space is provided. We find that cell types are not as clearly separated in the CPLVM and CGLVM shared latent spaces as they are for contrastiveVI.

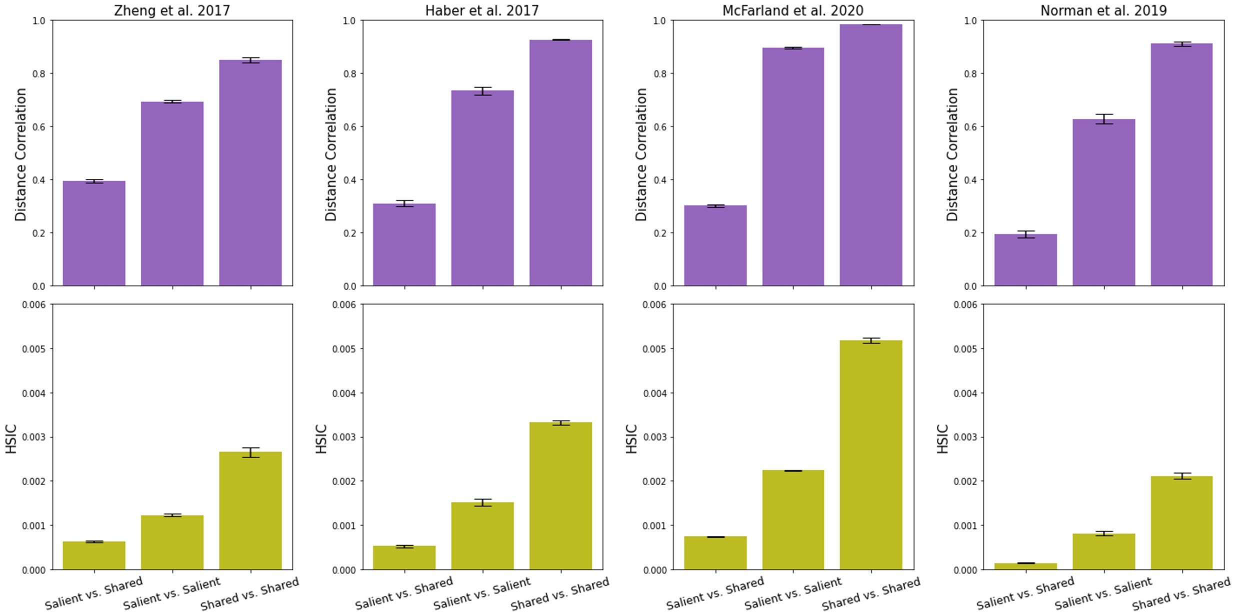

**Supplementary Figure 4: Evaluating disentanglement of contrastiveVI salient and shared latent space.** Mean and standard error of distance correlation and the HSIC (Hilbert-Schmidt Independence Criterion) estimate between contrastiveVI salient and shared latent spaces across five random model training trials are plotted. Distance correlation and the HSIC are dependence measures for multivariate random variables, with higher values indicating more statistical dependence (less disentanglement). Distance correlation and the HSIC estimate within the same latent space (i.e. comparison between *salient vs. salient* or *shared vs. shared*) are plotted as benchmarks for fully entangled latent variables. Results with salient and shared latent dimension = 10 are shown. See **Supplementary Note 7** for evaluation details.

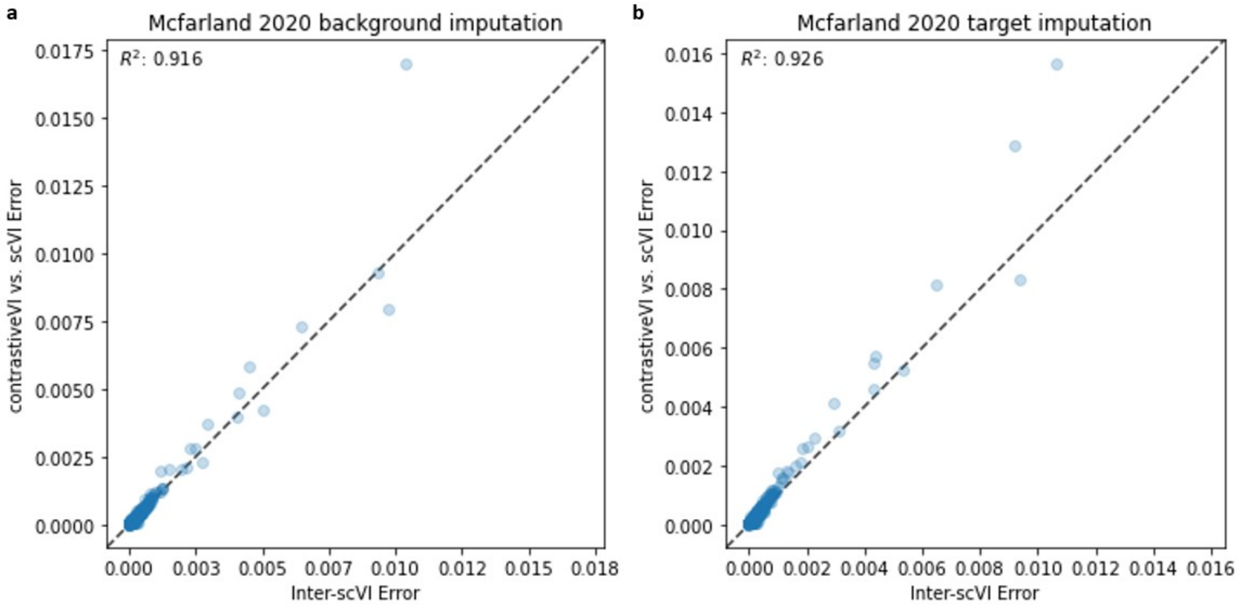

**Supplementary Figure 5: Evaluating the quality of contrastiveVI's imputed normalized expression values. a,** For each gene in the McFarland et al. [11] dataset, the mean absolute error was computed between imputed normalized expression values for background cells from an scVI model and a contrastiveVI model (y axis) or a second scVI model (x axis) trained with a different random initialization. We find strong agreement between the two sets of values. This agreement was quantified by computing the $R^2$ between the data and a line with intercept at zero and slope of one (i.e. the line $y = x$). **b,** The same procedure as in (**a**) was repeated for target cells. Once again we find the contrastiveVI-versus-scVI errors closely match the inter-scVI errors.

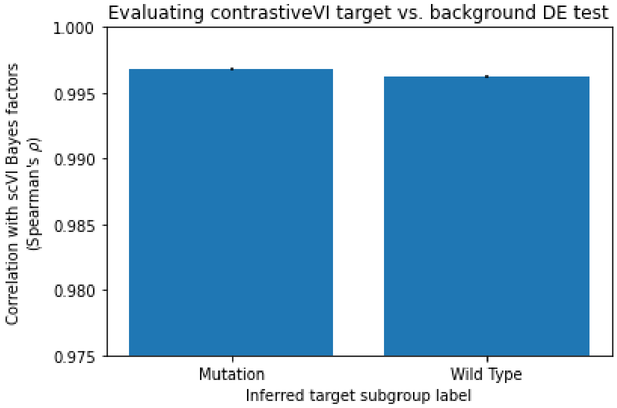

**Supplementary Figure 6: Evaluating the quality of contrastiveVI's target-versus-background DE test.** To assess the quality of contrastiveVI's target-versus-background DE test, we compared the resulting Bayes' factors from running the test on the two discovered target cell subgroups with those returned by an scVI model trained to perform the same task. In this case, the scVI model was trained on *all* cells (i.e., target and background). We find that the Bayes factors from contrastiveVI and scVI are highly correlated.

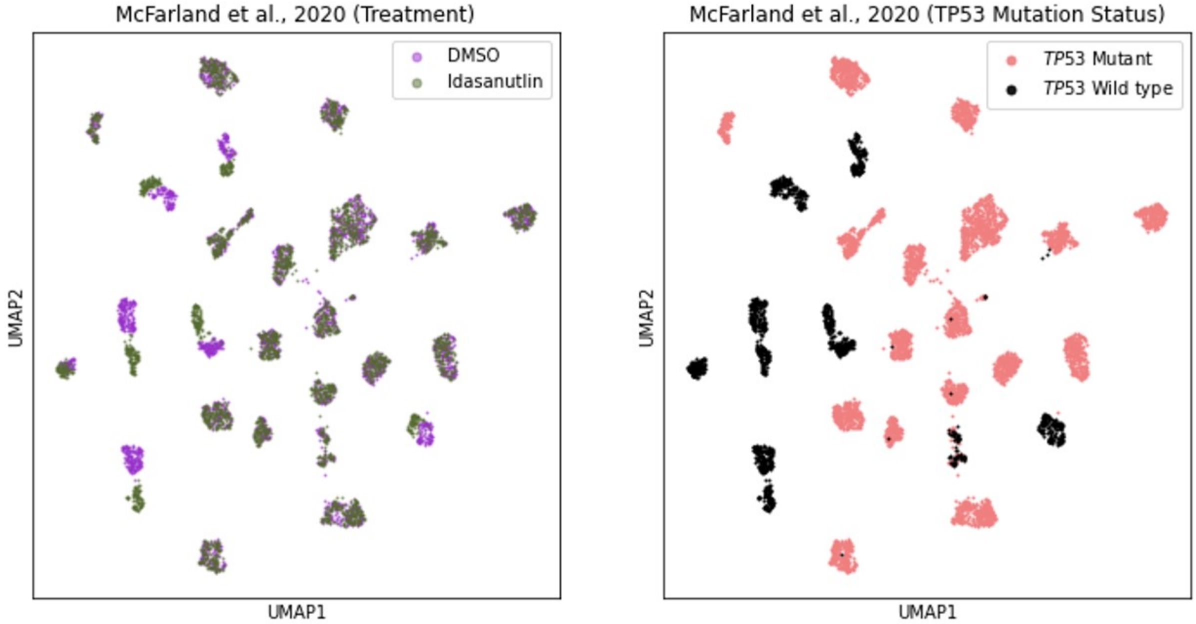

**Supplementary Figure 7: Cell line separation by treatment type in McFarland et al., 2020.** UMAP plots of library-size-normalized and log-transformed data from McFarland et al. [11] colored by treatment type (left) and *TP53* mutation status (right). Cells with wild type *TP53* clearly separate by treatment type.

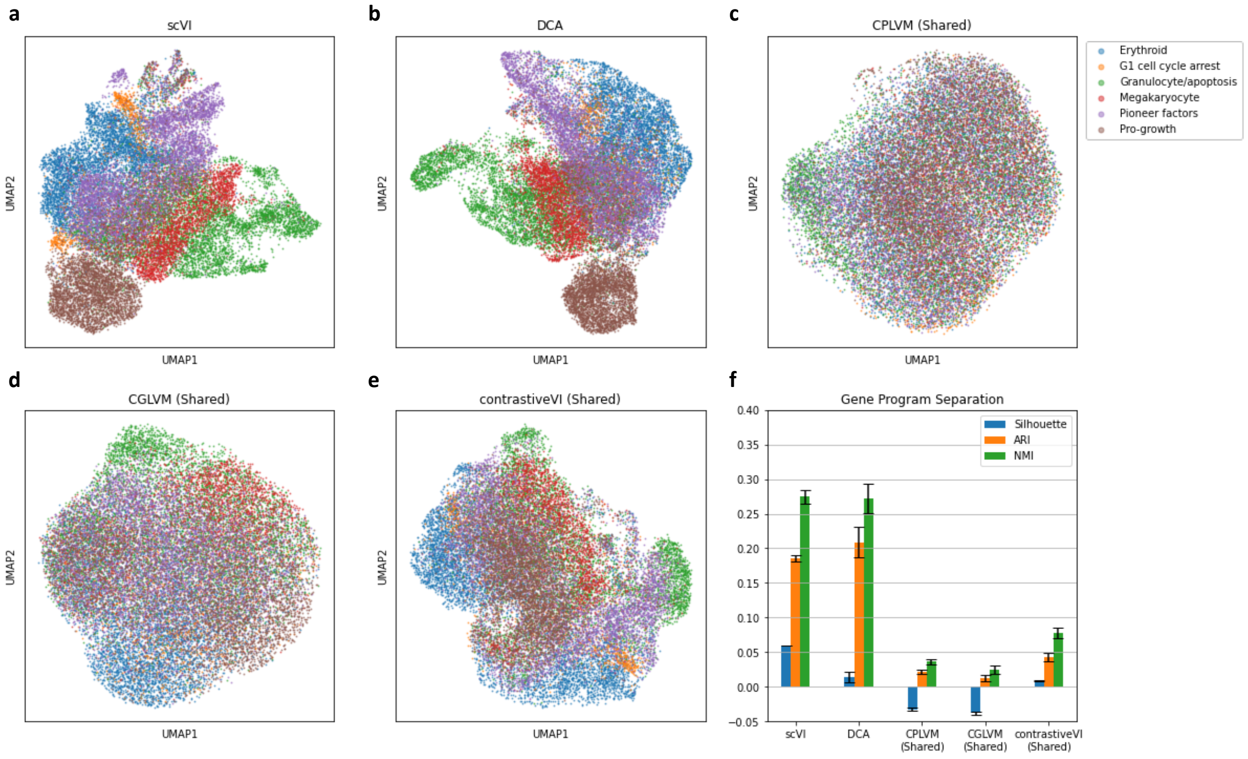

**Supplementary Figure 8: Shared variations captured by contrastiveVI and baseline models trained on Perturb-Seq data from Norman et al. [13]. a,b,c,d,e,** UMAP plots of shared latent spaces for contrastiveVI and baseline models. For scVI and DCA, we plot the model's single latent space. **f,** Metrics quantifying separation of cells by induced gene program in models' shared latent spaces. Here values closer to zero indicate better performance.

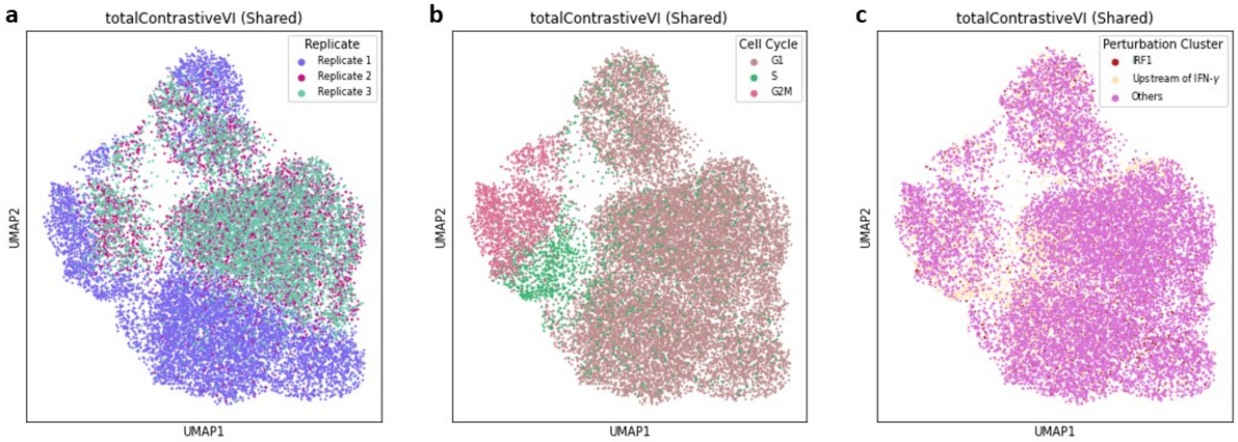

**Supplementary Figure 9: UMAP plots of the totalContrastiveVI shared latent space for Papalexi et al. [14]. a,b,c,** UMAP plots of the totalContrastiveVI shared latent space for Papalexi et al. [14] colored by replicate number (**a**), cell cycle stage (**b**) and perturbation cluster label as determined from the totalContrastiveVI salient latent space (**c**).

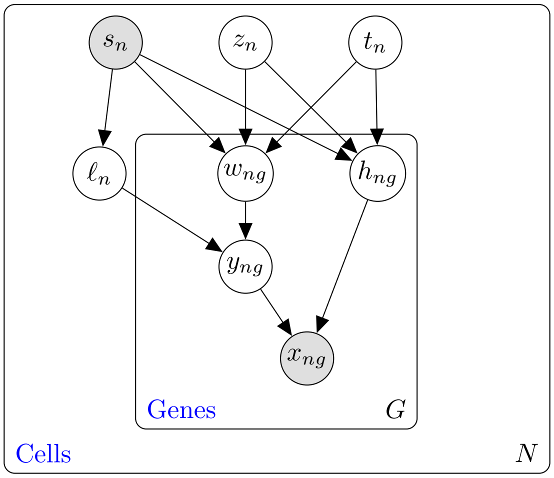

**Supplementary Figure 10: The contrastiveVI probabilistic graphical model.** Unshaded nodes represent latent variables, while shaded nodes represent observed variables. Edges denote conditional independence, while rectangles indicate independent replication.

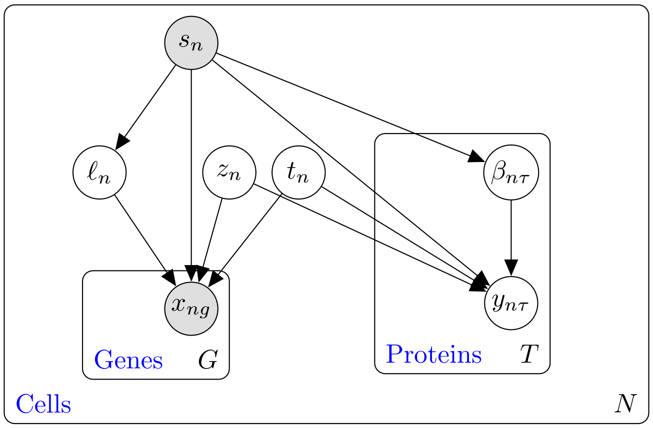

**Supplementary Figure 11: The totalContrastiveVI probabilistic graphical model.** Unshaded nodes represent latent variables, while shaded nodes represent observed variables. Edges denote conditional independence, while rectangles indicate independent replication.

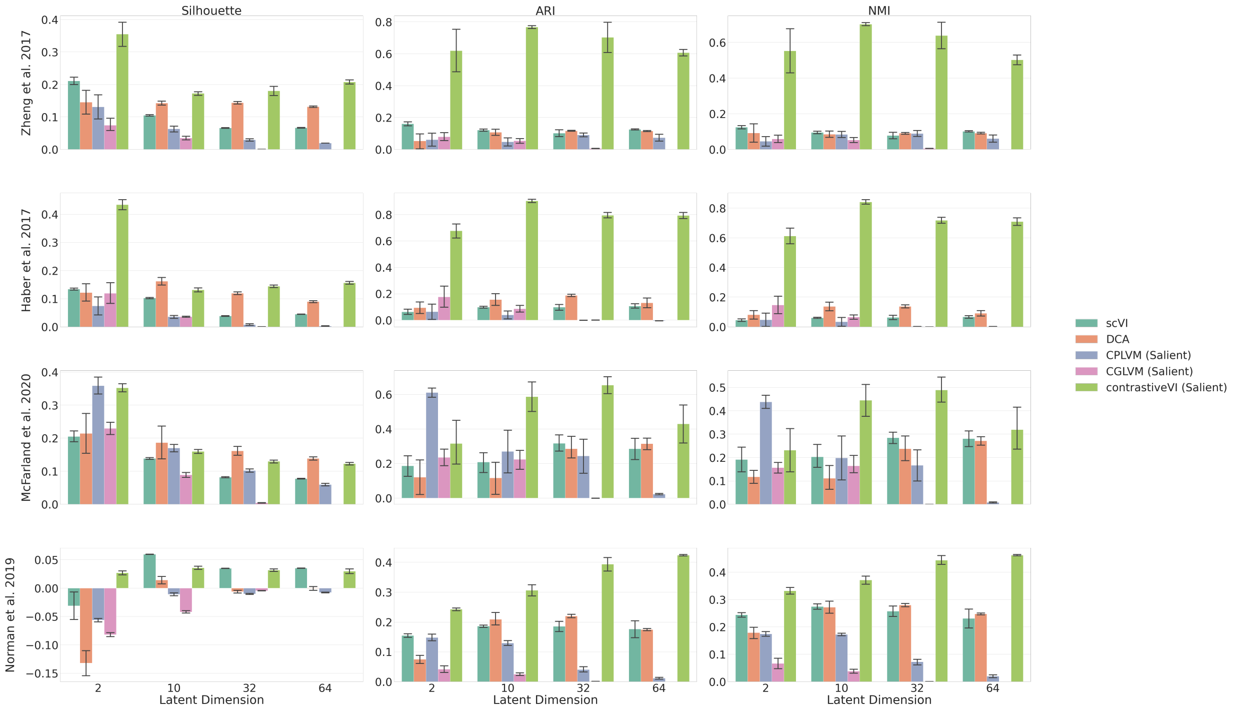

**Supplementary Figure 12: contrastiveVI performance with varying (salient) latent dimension.** Mean and standard error of average silhouette width (silhouette), adjusted Rand Index (ARI), and normalized mutual information (NMI) across five random model training trials are plotted for each method's (salient) latent variables at dimension $= 2, 10, 32, 64$ for all benchmark datasets. Results for CGLVM with dimension $= 64$ are not included due to numerical instabilities resulting in NaN values during optimization. Note y-axis scales vary in subplots.

### Supplementary Tables

| Dataset (Task in target dataset) | Model | Target subgroup separation Silhouette | ARI | NMI |
|---|---|---|---|---|
| Zheng 2017 (Pre- vs. post-transplant) | scVI | $0.105 \pm 0.002$ | $0.121 \pm 0.006$ | $0.095 \pm 0.007$ |
| | DCA | $0.143 \pm 0.007$ | $0.108 \pm 0.017$ | $0.085 \pm 0.017$ |
| | CPLVM | $0.063 \pm 0.009$ | $0.048 \pm 0.027$ | $0.084 \pm 0.018$ |
| | CGLVM | $0.035 \pm 0.005$ | $0.054 \pm 0.015$ | $0.053 \pm 0.014$ |
| | contrastiveVI | $\mathbf{0.173 \pm 0.005}$ | $\mathbf{0.768 \pm 0.009}$ | $\mathbf{0.702 \pm 0.008}$ |
| Haber 2017 (*Salmonella* vs. *H. poly*) | scVI | $0.102 \pm 0.002$ | $0.099 \pm 0.008$ | $0.061 \pm 0.004$ |
| | DCA | $\mathbf{0.162 \pm 0.012}$ | $0.156 \pm 0.040$ | $0.137 \pm 0.026$ |
| | CPLVM | $0.036 \pm 0.006$ | $0.039 \pm 0.027$ | $0.034 \pm 0.026$ |
| | CGLVM | $0.036 \pm 0.002$ | $0.086 \pm 0.024$ | $0.066 \pm 0.014$ |
| | contrastiveVI | $0.131 \pm 0.007$ | $\mathbf{0.904 \pm 0.012}$ | $\mathbf{0.841 \pm 0.015}$ |
| McFarland 2020 (*TP53* mutant vs wild-type) | scVI | $0.138 \pm 0.003$ | $0.208 \pm 0.066$ | $0.203 \pm 0.055$ |
| | DCA | $\mathbf{0.186 \pm 0.048}$ | $0.117 \pm 0.096$ | $0.112 \pm 0.047$ |
| | CPLVM | $0.170 \pm 0.010$ | $0.269 \pm 0.110$ | $0.198 \pm 0.084$ |
| | CGLVM | $0.089 \pm 0.007$ | $0.224 \pm 0.056$ | $0.165 \pm 0.035$ |
| | contrastiveVI | $0.161 \pm 0.003$ | $\mathbf{0.558 \pm 0.102}$ | $\mathbf{0.418 \pm 0.085}$ |
| Norman 2019 (Induced gene program) | scVI | $\mathbf{0.060 \pm 0.000}$ | $0.186 \pm 0.005$ | $0.274 \pm 0.010$ |
| | DCA | $0.014 \pm 0.006$ | $0.209 \pm 0.019$ | $0.272 \pm 0.019$ |
| | CPLVM | $-0.011 \pm 0.003$ | $0.130 \pm 0.008$ | $0.173 \pm 0.005$ |
| | CGLVM | $-0.042 \pm 0.002$ | $0.025 \pm 0.005$ | $0.039 \pm 0.007$ |
| | contrastiveVI | $0.036 \pm 0.003$ | $\mathbf{0.306 \pm 0.021}$ | $\mathbf{0.371 \pm 0.016}$ |

**Supplementary Table 1:** Quantitative evaluation of how well each method separates ground truth subclasses of target data points in the method's salient latent space (for CPLVM, CGLVM, and contrastiveVI) or the method's single latent space (for scVI and DCA).

| Pathway Name | Pathway Entry | Adjusted p-value |
|---|---|---|
| Hematopoietic cell lineage | hsa04640 | 9.35e-10 |
| Asthma | hsa05310 | 3.17e-08 |
| Systemic lupus erythematosus | hsa05322 | 4.91e-05 |
| Antigen processing and presentation | hsa04612 | 8.29e-05 |
| Type I diabetes mellitus | hsa04940 | 1.03e-04 |
| Allograft rejection | hsa05330 | 1.90e-04 |
| Graft-versus-host disease | hsa05332 | 3.41e-04 |
| Leishmaniasis | hsa05140 | 5.26e-04 |
| Cell adhesion molecules | hsa04514 | 5.26e-04 |
| Rheumatoid arthritis | hsa05323 | 1.09e-03 |
| Chagas disease | hsa05142 | 1.33e-03 |
| Toxoplasmosis | hsa05145 | 1.50e-03 |
| Staphylococcus aureus infection | hsa05150 | 3.18e-03 |
| Intestinal immune network for IgA production | hsa04672 | 4.04e-03 |
| NF-kappa B signaling pathway | hsa04064 | 4.04e-03 |
| Viral myocarditis | hsa05416 | 4.04e-03 |
| Tuberculosis | hsa05152 | 5.88e-03 |
| Autoimmune thyroid disease | hsa05320 | 7.14e-03 |
| Inflammatory bowel disease | hsa05321 | 7.61e-03 |
| Legionellosis | hsa05134 | 8.51e-03 |
| Influenza A | hsa05164 | 1.04e-02 |
| B cell receptor signaling pathway | hsa04662 | 1.59e-02 |
| VEGF signaling pathway | hsa04370 | 1.59e-02 |
| Glycine, serine and threonine metabolism | hsa00260 | 1.85e-02 |
| Cytokine-cytokine receptor interaction | hsa04060 | 1.89e-02 |
| HTLV-I infection | hsa05166 | 2.76e-02 |
| Transcriptional misregulation in cancer | hsa05202 | 2.76e-02 |
| Fc epsilon RI signaling pathway | hsa04664 | 2.81e-02 |
| Apoptosis | hsa04210 | 3.48e-02 |
| Primary immunodeficiency | hsa05340 | 4.57e-02 |
| Pertussis | hsa05133 | 4.57e-02 |
| Colorectal cancer | hsa05210 | 4.57e-02 |
| Arachidonic acid metabolism | hsa00590 | 4.57e-02 |
| Osteoclast differentiation | hsa04380 | 4.57e-02 |
| Arginine and proline metabolism | hsa00330 | 4.63e-02 |
| T cell receptor signaling pathway | hsa04660 | 4.69e-02 |

**Supplementary Table 2:** All pathways found to be enriched (false discovery rate $< 0.05$) based on the differentially expressed genes for the two clusters in contrastiveVI's salient latent space for the dataset collected by Zheng et al., 2017.

| Pathway Name (Associated Differentially Expressed Genes) | Pathway Entry | Adjusted p-value |
|---|---|---|
| Cell cycle *Homo sapiens* (*CCNA2*, *CDKN1A*, *PLK1*, *MDM2*) | hsa04110 | 1.66e-2 |
| p53 signaling pathway *Homo sapiens* (*CDKN1A*, *TP53I3*, *MDM2*) | hsa04115 | 2.14e-2 |
| Viral carcinogenesis *Homo sapiens* (*CCNA2*, *CDKN1A*, *MDM2*, *HIST1H4C*) | hsa05203 | 3.61e-2 |

**Supplementary Table 3:** All pathways found to be enriched (false discovery rate $< 0.05$) based on the differentially expressed genes for the *TP53* wild type cluster found in contrastiveVI's salient latent space versus background cells for the dataset collected in [11]

| Model | Model Characteristics | | | | | | Software Capabilities | | |
|---|---|---|---|---|---|---|---|---|---|
| | Expressive | Count Distribution | Over-dispersion | Library size | Multimodal extensions | Generative Model | Dimension Reduction | Differential Expression | Imputation |
| mixseq | | | | | | | ✓ | | |
| cPCA | | | | | | | ✓ | | |
| PCPCA | | | | | | | ✓ | | ✓ |
| CLVM | | | | | | | ✓ | | |
| CPLVM | | ✓ | | ✓ | | ✓ | ✓ | ✓ | |
| CGLVM | | ✓ | | ✓ | | ✓ | ✓ | ✓ | |
| cVAE | ✓ | | | | | ✓ | ✓ | | ✓ |
| contrastiveVI | ✓ | ✓ | ✓ | ✓ | ✓ | ✓ | ✓ | ✓ | ✓ |

**Supplementary Table 4:** Comparison of contrastiveVI with previous methods for contrastive analysis. We compare across both features of the models as well as capabilities of their corresponding software packages. *Expressive*: Indicates whether the model can capture nonlinear relationships. *Count Distribution*: Denotes whether the distribution modeling the data has support in the integer set. *Over-dispersion*: Indicates whether the count-based distribution modeling the data accounts for variance being greater than the mean. *Library size*: Denotes whether the model corrects for library size differences. *Batch effects*: Whether the model can account for nuisance variations due to differences in experimental conditions. *Generative Model*: Indicates if the model supports sampling from a distribution.

| Dataset | Num. Samples (background) | Num. Samples (target) | Platform |
|---|---|---|---|
| Zheng et al. 2017 | 4,457 | 12,399 | GemCode |
| Haber et al., 2017 | 3,240 | 4,481 | SMART-Seq2 |
| McFarland et al., 2020 | 2,831 | 3,097 | MIX-Seq |
| Norman et al., 2019 | 8,907 | 24,913 | Perturb-Seq |
| Papalexi et al., 2021 | 2,386 | 18,343 | ECCITE-Seq |

**Supplementary Table 5:** Summary of datasets used.

| Cell Type | Number of cells | | |
|---|---|---|---|
| | Healthy (background) | *Salmonella* (target) | *H. polygyrus* (target) |
| Endocrine | 112 | 69 | 82 |
| Enterocyte | 424 | 705 | 128 |
| Enterocyte.Progenitor | 545 | 229 | 586 |
| Goblet | 216 | 126 | 317 |
| Stem | 670 | 207 | 592 |
| TA | 421 | 112 | 353 |
| TA.Early | 792 | 300 | 436 |
| Tuft | 60 | 22 | 217 |

**Supplementary Table 6:** Number of cell types present in each condition for the dataset by Haber et al. [5].

| Cell Line | Number of cells | |
| --- | --- | --- |
| | DMSO-treated (background) | Idasanutlin-treated (target) |
| BICR6_UPPER_AERODIGESTIVE_TRACT | 82 | 111 |
| BICR31_UPPER_AERODIGESTIVE_TRACT | 245 | 277 |
| BT474_BREAST | 53 | 71 |
| BT549_BREAST | 100 | 131 |
| CAOV3_OVARY | 97 | 140 |
| CCFSTTG1_CENTRAL_NERVOUS_SYSTEM | 103 | 77 |
| COLO680N_OESOPHAGUS | 129 | 129 |
| COV434_OVARY | 60 | 75 |
| DKMG_CENTRAL_NERVOUS_SYSTEM | 103 | 93 |
| IALM_LUNG | 105 | 141 |
| LNCAPCLONEFGC_PROSTATE | 139 | 113 |
| LS1034_LARGE_INTESTINE | 72 | 118 |
| NCIH226_LUNG | 165 | 94 |
| NCIH2347_LUNG | 111 | 159 |
| RCC10RGB_KIDNEY | 172 | 114 |
| RCM1_LARGE_INTESTINE | 109 | 133 |
| RERFLCAD1_LUNG | 99 | 123 |
| SH10TC_STOMACH | 123 | 122 |
| SKMEL2_SKIN | 150 | 141 |
| SKMEL3_SKIN | 145 | 183 |
| SNU1079_BILIARY_TRACT | 101 | 105 |
| SQ1_LUNG | 113 | 150 |
| TEN_ENDOMETRIUM | 155 | 177 |
| UMUC1_URINARY_TRACT | 100 | 120 |

**Supplementary Table 7:** Number of cells by cell line present in each condition for the dataset by McFarland et al. [11].

### Supplementary Note 1

#### Further discussion on the relationship between contrastive analysis and batch effect correction

Here we provide more details on the relationship between batch effect correction and contrastive analysis. A batch effect refers to systematic variations between data generated at separate time points ("batches") from technical factors unrelated to any underlying biological phenomena. For example, batch effects may be present in single-cell datasets due to variations in cell dissociation and handling protocols, library preparation strategies, or differences between sequencing platforms [6]. When data originating from multiple batches are analyzed, batch effects may lead to spurious conclusions; as a result, many methods have been developed to correct for batch effects in single-cell data while preserving true biological signals in the data [6, 16, 10].

On the other hand, in the contrastive analysis setting we assume that the variations enriched in a target dataset compared to the background are biologically meaningful and worthy of further study. Thus, rather than *removing* these enriched variations—as would be done by applying a batch effect correction method—we instead seek to *isolate* them for further analyses. By doing so, target-specific heterogeneity can be explored to better understand the effect of a given treatment on the target cells.

The reader may wonder: do batch effects impact the performance of contrastive analysis methods? In practice, as shown by our experiments in the main text, we found that variations in the contrastiveVI salient space consistently captured known biological phenomena as opposed to technical factors. Indeed, for datasets that we may wish to analyze with contrastive analysis, their experimental designs often preclude the potential issue of batch effects. For example, in many cases, an entire dataset (including target and background cells) is collected in a single batch; this was the case for the Perturb-Seq dataset from Norman et al. [13] analyzed in this study. In other cases, the data are collected in two batches, which correspond exactly to target versus background. For example, in the dataset from McFarland et al. [11] analyzed in this study, the background (cells treated with a DMSO control solution) and target (cells treated with idasanutlin) were measured separately. Nevertheless, because *only* target cells are embedded in the salient latent space and all of the target cells were collected in a single batch, batch effects should not confound our analyses. Finally, even when batch effects were present, as in the ECCITE-Seq dataset analyzed in this study [14], we found that the batch-effect-induced variations were relegated to contrastiveVI's shared latent space while the salient latent space was not affected.

### Supplementary Note 2

#### Further discussion on the relationship between contrastive analysis and differential abundance testing

Here we expand upon the differences between contrastive analysis and differential abundance testing. Methods for differential abundance testing, such as MELD [1], MILO [2], and DASeq

[18], aim to identify subpopulations of cells whose distributions differ between two or more predefined biological states. However, once these populations are identified, such methods do not attempt to disentangle variations in these cells that are shared across conditions from those that are specific to one condition. On the other hand, methods for contrative analysis explicitly decompose variations in the data into shared and target-specific latent factors. Recovering these target-specific latent factors may assist in identifying relationships between different subpopulations of cells in the target condition that are not obvious in the original gene expression space.

For example, consider the MIX-Seq dataset from McFarland et al. [11] analyzed in this study. This dataset measured cancer cell lines' transcriptional responses to idasanutlin. It is well known that cell lines' response to this drug is governed by the *TP53* gene. That is, if a cell line has wild type *TP53*, it will experience activation of the p53 pathway when exposed to idasanutlin; on the other hand, cell lines with transcriptionally inactive mutant *TP53* will not respond to the drug. Differential abundance testing methods may identify the cells that were affected by idasanutlin exposure. However, due to differences between cell lines that are unrelated to idasanutlin, differential abundance testing methods would likely identify multiple distinct subpopulations of perturbed cells. Such a result could obscure the fact that perturbed cells' responses were the result of a single shared biological mechanism. On the other hand, as demonstrated in the main text, we find that contrastiveVI's salient latent space successfully isolates this shared mechanism underlying cell lines' responses to idasanutlin.

### Supplementary Note 3

#### Integrating out contrastiveVI's latent variables

We first show that if

$$\begin{aligned} w &\sim \text{Gamma}(\rho, \theta) \\ y|w &\sim \text{Poisson}(\ell w), \end{aligned}$$

where $\rho, \theta \in \mathbb{R}_+$ are the mean and shape parameter of the gamma distribution, respectively, and $\ell \in \mathbb{R}_+$, then $y$ follows a negative binomial distribution:

$$
\begin{aligned}
p(y) &= \int p(y|w)p(w)dw \\
&= \int \frac{\ell^y w^y e^{-\ell w}}{\Gamma(y+1)} \frac{\left(\frac{\theta}{\rho}\right)^\theta w^{\theta-1} e^{-\theta w/\rho}}{\Gamma(\theta)} dw \\
&= \frac{\ell^y \left(\frac{\theta}{\rho}\right)^\theta}{\Gamma(y+1)\Gamma(\theta)} \int w^{y+\theta-1} e^{-\left(\ell+\frac{\theta}{\rho}\right)w} dw \\
&= \frac{\ell^y \left(\frac{\theta}{\rho}\right)^\theta}{\Gamma(y+1)\Gamma(\theta)} \frac{\Gamma(y+\theta)}{\left(\ell+\frac{\theta}{\rho}\right)^{y+\theta}} \\
&= \frac{\Gamma(y+\theta)}{\Gamma(y+1)\Gamma(\theta)} \left(\frac{\theta}{\ell\rho+\theta}\right)^\theta \left(\frac{\ell\rho}{\ell\rho+\theta}\right)^y.
\end{aligned}
$$

The integral in the third line is evaluated by observing that the integrand is the unnormalized probability density function of a gamma distribution. The final line is exactly the probability mass function of a negative binomial distribution with mean $\ell\rho$ and inverse dispersion $\theta$.

Next, we can incorporate multiplication of $y$ by zero as a mixture between a point mass at zero and the original distribution of $y$. This enables us to write the probability mass function of $p(x_{ng}|z_n, t_n, \ell_n, s_n)$ as

$$
\begin{cases}
p(x_{ng} = 0|v_n, \ell_n) = f_h^g(v_n) + (1 - f_h^g(v_n)) \left(\dfrac{\theta_g}{\ell_n f_w^g(v_n) + \theta_g}\right)^{\theta_g} \\
p(x_{ng} = y|v_n, \ell_n) = (1 - f_h^g(v_n)) \dfrac{\Gamma(y+\theta_g)}{\Gamma(y+1)\Gamma(\theta_g)} \left(\dfrac{\theta_g}{\ell_n f_w^g(v_n) + \theta_g}\right)^{\theta_g} \left(\dfrac{\ell_n f_w^g(v_n)}{\ell_n f_w^g(v_n) + \theta_g}\right)^y,
\end{cases}
$$

where $v_n = \{z_n, t_n, s_n\}$ and $y \in \mathbb{N}^+$. Letting $f_w$ encode the mean of $w$ and $f_h$ the probability of technical dropout, this is exactly the probability mass function of a zero-inflated negative binomial (ZINB) distribution.

### Supplementary Note 4

#### Deriving contrastiveVI's evidence lower bounds

Here, we derive the variational lower bounds for contrastiveVI presented in the main text. For a given target cell $x$, the contrastiveVI generative model's joint likelihood function factorizes as

$$p(x, z, t, \ell | s) = p(x | z, t, \ell, s) p(\ell | s) p(z) p(t).$$

Next, in order to perform variational inference we define the variational posterior as

$$q(z, t, \ell | x, s) = q(z | x, s) q(t | x, s) q(\ell | x, s).$$

Then we have

$$\begin{aligned}
\log p(x|s) &= \log \int p(x, z, t, \ell | s) dz dt d\ell \\
&= \log \int \frac{p(x, z, t, \ell | s) q(z, t, \ell | x, s)}{q(z, t, \ell | x, s)} dz dt d\ell \\
&\geq \int q(z, t, \ell | x, s) \log \frac{p(x, z, t, \ell | s)}{q(z, t, \ell | x, s)} dz dt d\ell \\
&= \int q(z, t, \ell | x, s) \log \frac{p(x | z, t, \ell, s) p(z, t, \ell | s)}{q(z, t, \ell | x, s)} dz dt d\ell \\
&= \int \left( q(z, t, \ell | x, s) \log p(x | z, t, \ell, s) + q(z, t, \ell | x, s) \log \frac{p(z, t, \ell | s)}{q(z, t, \ell | x, s)} \right) dz dt d\ell \\
&= \mathbb{E}_{q(z,t,\ell|x,s)}[\log p(x | z, t, \ell, s)] - D_{KL}(q(z, t, \ell | x, s) \,||\, p(z, t, \ell | s)) \\
&= \mathbb{E}_{q(z,t,\ell|x,s)}[\log p(x | z, t, \ell, s)] - D_{KL}(q(z | x, s) \,||\, p(z)) \\
&\quad - D_{KL}(q(t | x, s) \,||\, p(t)) - D_{KL}(q(\ell | x, s) \,||\, p(\ell | s)),
\end{aligned}$$

where we use Jensen's inequality in the third step and the independence of $z$, $t$, and $\ell$ to decompose the KL divergence term in the last step. Next, for a background point $b$, we assume our generative process factorizes as

$$p(b, z, \ell | s) = p(b | z, \ell, s) p(\ell | s) p(z),$$

with a corresponding variational posterior of

$$q(z, \ell | b, s) = q(z | b, s) q(\ell | b, s).$$

We then have

$$
\begin{aligned}
\log p(b|s) &= \log \int p(b, z, \ell|s) dz d\ell \\
&= \log \int \frac{p(b, z, t, \ell|s) q(z, \ell|b, s)}{q(z, \ell|b, s)} dz d\ell \\
&\geq \int q(z, \ell|b, s) \log \frac{p(b, z, \ell|s)}{q(z, \ell|b, s)} dz d\ell \\
&= \int q(z, \ell|b, s) \log \frac{p(b|z, \ell, s) p(z, \ell|s)}{q(z, \ell|b, s)} dz d\ell \\
&= \int \left( q(z, \ell|b, s) \log p(b|z, \ell, s) + q(z, \ell|b, s) \log \frac{p(z, \ell|s)}{q(z, \ell|b, s)} \right) dz d\ell \\
&= \mathbb{E}_{q(z,\ell|b,s)}[\log p(b|z, \ell, s)] - D_{KL}(q(z, \ell|b, s) \,||\, p(z, \ell|s)) \\
&= \mathbb{E}_{q(z,\ell|b,s)}[\log p(b|z, \ell, s)] - D_{KL}(q(z, |b, s) \,||\, p(z)) - D_{KL}(q(\ell, |x, s) \,||\, p(\ell|s)).
\end{aligned}
$$

### Supplementary Note 5

#### Further details on contrastiveVI network architecture

Three separate encoder neural networks were used to parameterize our approximate posterior distributions for $z$, $t$, and $\ell$. Each network had a single hidden layer consisting of 128 nodes. This was followed by a batch normalization layer [7], a rectified linear unit (ReLU) activation function [12], and a dropout layer [15]. During training, the dropout probability was set to 10%. The resulting 128 node values were then used as inputs for two linear layers that parameterized the given factor (e.g., for the encoder corresponding to $q(z|x, s)$, the linear layers parameterized the mean and variance of $z$). For our main results, we used 10-dimensional mean and variance parameters for $z$ and $t$, and we used a 1-d mean and shape parameter for $\ell$.

Our decoder network began with a single hidden layer taking in values of our three latent factors (i.e., $z$, $t$ and $\ell$) with an output dimension of 128. This was followed by batch normalization, a ReLU activation function, and a dropout layer as described previously. The output of this sequence was then fed to three separate decoder layers, one for each of the three parameters of the ZINB distribution. To force the ZINB scale parameter to lie between 0 and 1, we applied a softmax activation function to its corresponding decoder's output. We note that similar decoding approaches have been successfully used by previous unsupervised modeling approaches for scRNA-seq data [10, 3].

### Supplementary Note 6

#### Further details on totalContrastiveVI network architecture

To parameterize $z, t$, and $\ell$, encoder networks with the same architecture as those in contrastiveVI were used. To parameterize $q(\beta|z, t, s)$, we used a neural network with one hidden layer of 128 nodes that takes in as input $(z, t, s)$ and outputs the parameters of $q(\beta|z, t, s)$. As in the other encoder networks, the hidden nodes were followed by a batch normalization layer, a ReLU activation function, and a dropout layer with dropout probability set to 10%.

The decoder consisted of three individual neural networks with one hidden layer of 128 nodes. Each network took as input our latent factors $z$ and $t$ as well as covariate labels $s$. The first network mapped to the parameters of the mean of the RNA likelihood $\rho_n$. The second network mapped to the foreground mean of the protein likelihood $\alpha_n$. The third network mapped to the mixing parameter $\pi_n$ of the protein likelihood mixture. All networks used batch normalization, a ReLU activation function in the hidden layer, and a dropout layer as described previously. To force $\pi_n$ to lie between zero and one, an additional sigmoid activation function was applied to the output of its network. We note that our architecture closely follows the default totalVI architecture as implemented in `scvi-tools` with the addition of the salient variables $t$ and corresponding encoder for $t$ as well as some minor differences in hyperparameter choices (e.g. 128 hidden nodes per layer in our architecture as compared to 256 in totalVI).

### Supplementary Note 7

#### Details on disentanglement evaluation.

We applied distance correlation [17] and the HSIC (Hilbert-Schmidt Independence Criterion) [4], which are measures of statistical dependence between two multivariate random variables, to quantitatively evaluate the disentanglement between contrastiveVI's salient latent space and shared latent space. Particularly, when salient latent variables and shared latent variables are more disentangled, they are less statistically dependent and hence have lower dependence measures. Distance correlation and the HSIC are valid disentanglement metrics for multivariate random variables and have been used as objectives to achieve disentangled representation learning [8, 9, 10].

For each dataset, samples of salient and shared latent variables were drawn from the contrastiveVI variational posterior distributions, conditioned on the *same* cells in the target dataset, to calculate distance correlation and the HSIC value. The radial basis function kernel with the free parameter $\sigma = 1$ was used for the HSIC. To construct mean estimates of distance correlation and the HSIC, this Monte Carlo sampling procedure was repeated 100 times for each dataset. Due to GPU memory constraints, a different random subset of 20,000 target cells was used each time to draw Monte Carlo samples from the Norman et al. target dataset (total number of target cells is 24,913). To interpret the dependence measure values, we consider two sets of samples drawn from the *same* latent space (i.e. salient or shared latent space) conditioned on the *same* target cells as a benchmark of entangled random variables. That is, if contrastiveVI's salient and shared latent variables are indeed disentangled, samples from the salient latent space and samples from the shared latent space should have lower dependence measures than samples drawn from only the salient or only the shared latent space.
